# *Magel2* deficiency drives microglial synaptic engulfment and alters neuron-microglia interactions in Prader-Willi syndrome

**DOI:** 10.64898/2026.09.30.755583

**Authors:** Jemima Helen, Moritz C. Wimmer, Berkan Kanmaz, Tim Schubert, Ingrid Possa Paranhos, Felipe Correa-da-Silva, Yuval Podpecan, Aliyu Jabir Muhammad, Eva M. Eisemann, Nadia Illma, Derya S. Arslan, Megan Iminitoff, Franziska Schommer, Henning Fröhlich, Felix Franke, Zuguang Gu, Daniel Hübschmann, Marnie E. Blewitt, Hui Yu, Malcolm J. Low, Vanessa Vogel, Daniel Kazdal, Elena G. Bochukova, Matthew K. Kirchner, Tim Gruber, I. Sadaf Farooqi, Francoise Muscatelli, Dick Swaab, Chun-Xia Yi, Han L. Tan, Ferdinand Althammer

**Affiliations:** Institute of Human Genetics, Heidelberg University Hospital, Heidelberg, Germany; Netherlands Institute for Neuroscience, Royal Netherlands Academy of Arts and Sciences (KNAW), Amsterdam, The Netherlands; Laboratory of Molecular Genetics, Howard Hughes Medical Institute, The Rockefeller University, New York, NY, USA; The Walter and Eliza Hall Institute of Medical Research, 1G Royal Parade, Parkville, Melbourne, Australia; Department of Medical Biology, University of Melbourne, Melbourne, Australia; Institute of Cell and Gene Technology, Shenzhen University of Advanced Technology, Shenzhen, China; Bioinformatics and Omics Data Analytics, DKFZ, Heidelberg, Germany; National Center for Tumor Diseases (NCT) Heidelberg; Department of Animal Sciences, The Ohio State University, Columbus, OH, United States; Department of Molecular and Integrative Physiology, University of Michigan Medical School, United States; Institute of Pathology, Heidelberg University Hospital, Heidelberg, Germany; Translational Lung Research Center (TLRC) Heidelberg, member of the DZL, Heidelberg, Germany; Blizard Institute, Barts and The London School of Medicine and Dentistry, Queen Mary University of London, London, UK; Department of Medical Physiology, College of Medicine, Texas A&M University, Bryan, TX, USA; Institute for Translational Metabolism Research, Helmholtz Diabetes Center at Helmholtz Munich, Neuherberg, Germany; German Center for Diabetes Research (DZD), 85764 Neuherberg, Germany; University of Cambridge, Metabolic Research Laboratories and NIHR Cambridge Biomedical Research Centre, Addenbrookes Hospital, Cambridge, United Kingdom; Institut de Neurobiologie de la Méditerranée (INMED), INSERM, Aix-Marseille Université, 13273 Marseille, France; Department of Anatomy and Neurobiology, University of California, Irvine, Irvine, CA 92617, USA

## Abstract

Prader–Willi syndrome (PWS) is a neurodevelopmental disorder characterized by hypothalamic dysfunction, developmental delay, hyperphagia, and cognitive impairment, yet the cellular mechanisms underlying circuit abnormalities remain unclear. Here, we show that loss of the imprinted gene Magel2 remodels microglial state and function in a mouse model of PWS. Three-dimensional morphometric analysis of hypothalamic microglia revealed reduced structural complexity and contracted territorial domains, emerging early in development and largely independent of obesity. Single-nucleus RNA sequencing of hypothalamic brain tissue identified extensive transcriptional reprogramming involving immune signaling, cellular communication, ribosome function, synaptic organization, and protein translation, with stronger effects in females. Magel2 deficiency also increased microglia-neuron contacts and synaptic engulfment, accompanied by reduced synaptic marker density, suggesting aberrant refinement of hypothalamic circuits. Analyses of human post-mortem hypothalamic tissue from individuals with PWS confirmed microglial structural alterations and increased complement engagement. Ligand-receptor inference further revealed reorganization of microglia-centered communication networks, including altered adhesion, trophic, and immune signaling consistent with enhanced phagocytic states. Together, these findings identify microglial remodeling and increased synaptic engagement as central features of Magel2 deficiency, implicate microglia in hypothalamic circuit dysfunction in PWS, and highlight excessive synaptic pruning as a novel dysfunctional mechanism of PWS.

## Introduction

Prader-Willi syndrome (PWS) is a rare genetic disorder caused by deletion of paternally inherited genes and non-coding RNAs on chromosome 15q11–q13 or maternal uniparental disomy of chromosome 15. Phenotypes commence with infantile hypotonia, feeding difficulties and failure to thrive [1]. Subsequently, individuals develop hyperphagia, characterized by excessive eating and early-onset severe obesity. Hypogonadism resulting in genital hypoplasia and infertility is common in affected individuals [2–4]. Among the genes affected in the PWS locus, *MAGEL2* has emerged as a key regulator of hypothalamic function [5], with roles in intracellular trafficking, neuropeptide processing [6], and energy homeostasis, thereby contributing to core features of PWS symptomatology [7]. Schaaf-Yang syndrome, caused by truncating variants in *MAGEL2*, shares several clinical features with PWS, including hypotonia, developmental delay, and endocrine abnormalities [8–9]. Hypothalamic transcriptome studies performed in *Magel2* knockout (KO) mice, a well-established murine model for PWS, report upregulation of microglia activation markers such as *Cxcl10* and *Tlr2* as well as increased expression of inflammatory markers in the hypothalamus [10]. In addition, bulk RNA sequencing of post-mortem hypothalamic tissue from PWS patients displayed upregulation of microglial genes such as *Ccl2*, *Il-1*β and *Cd14,* involved in inflammatory and innate immune response [11]. The data also showed differential gene expression in astrocytes and oligodendrocytes as well as downregulation of neuronal growth and developmental genes, highlighting glial- and neuron-specific transcriptomic changes suggesting their dysfunction in PWS [11]. A recent study found that microglia display fragmented processes, reduced branching, defective phagolysosome activity and reduced lysosomal marker expression in post-mortem hypothalamic tissue of PWS patients [12]. Microglia, the resident immune cells of the central nervous system, are increasingly recognized as key regulators of brain homeostasis and as critical contributors to a wide range of neurological and neurodevelopmental disorders [13–14]. Microglia are highly dynamic and continuously survey the neural environment through motile processes [15]. Beyond their immune functions, microglia form specialized contacts with neurons that enable activity-dependent monitoring and modulation of neuronal function, thereby contributing to synaptic regulation and circuit homeostasis [15–19].

Despite accumulating evidence implicating microglial dysfunction in PWS, it remains unclear whether these alterations are cell-intrinsic consequences of *Magel2* deficiency and how they impact microglia-neuron interactions during critical developmental windows. In particular, prior studies have largely relied on bulk transcriptomic approaches or static observations, leaving the structural and functional properties of microglia insufficiently characterized at single-cell resolution. Here, we combined high-resolution three-dimensional morphological analysis, single-nucleus (sn) transcriptomics utilizing novel tools for microglial state assessment, and computational inference of intercellular communication to systematically define microglial states in *Magel2* KO mice. By integrating these approaches with human post-mortem validation, we aimed to determine how Magel2 loss reshapes microglial function and contributes to hypothalamic circuit abnormalities in PWS, and to identify microglia-mediated dysfunctional synaptic pruning as a novel hallmark of the disorder.

## Results

To gain unbiased insight into potential alterations across different cell types in the *Magel2* KO mouse, we performed snRNA-seq analysis of hypothalamic tissue from *Magel2* KO animals and wild-type (WT) littermates. We chose P4, a developmental stage at which microglia become actively involved in synaptic refinement and developmental pruning [20] and where first functionally distinct microglial states emerge [21–22]. The overall analytical workflow is shown in **Figure 1A**. Integration and clustering of the datasets identified the major neural and glial populations in both sexes, including excitatory and inhibitory neurons, astrocytes, oligodendrocyte lineage cells, and microglia (**Figure 1B, C**). To validate the cell-type annotation, canonical marker gene expression was visualized across the identified cell populations and genotypes. The marker heatmaps showed selective enrichment of expected neuronal, glial, vascular, immune, and cycling-cell markers within their corresponding annotated clusters in both female and male datasets, supporting the reliability of the annotation strategy used for downstream analyses (**Supplementary Figure 1A, B**). The overall cellular composition was comparable between WT and *Magel2* KO samples, allowing direct assessment of genotype-dependent transcriptional changes within microglia.

**Figure 1.**
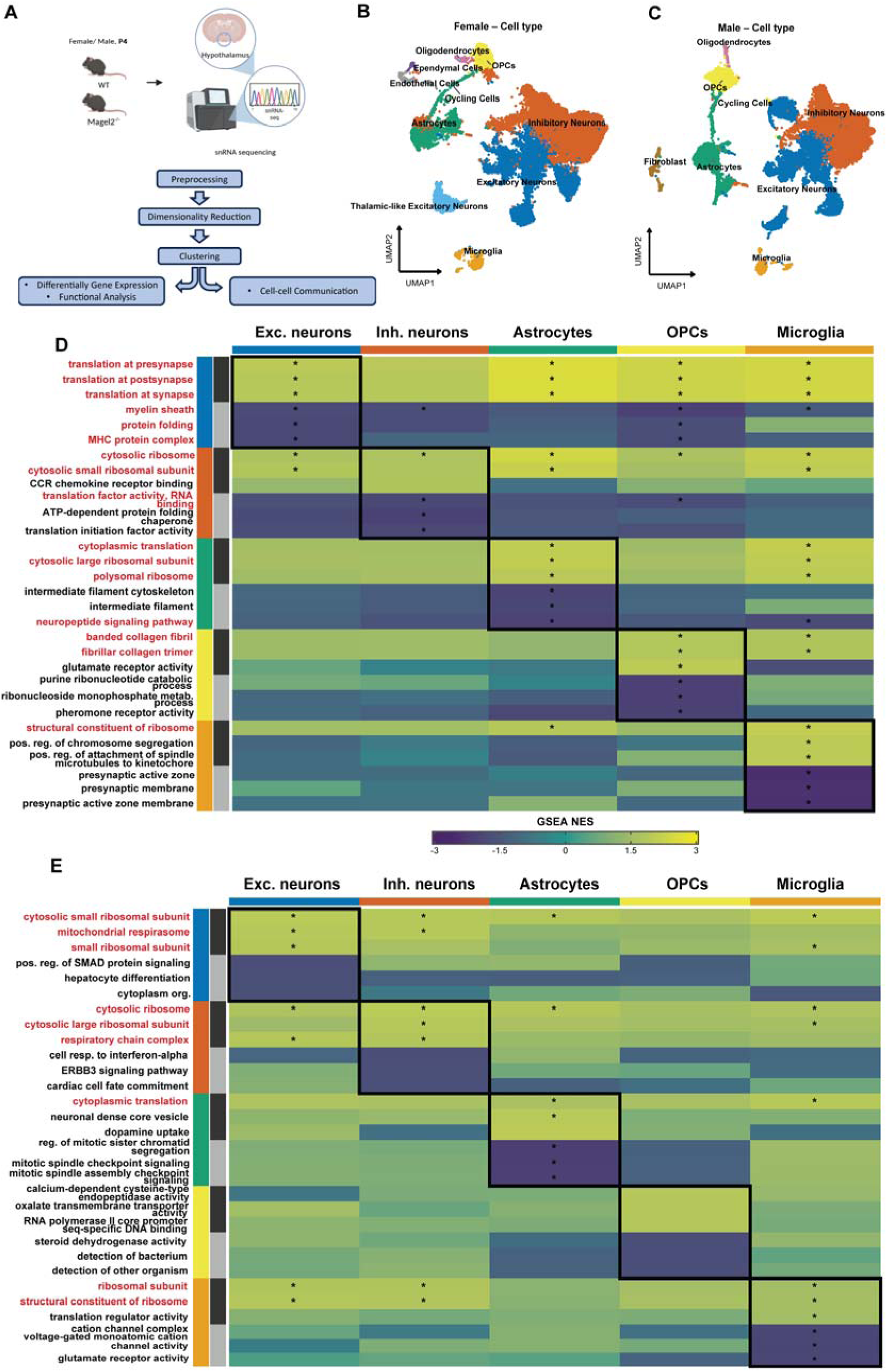
Single-nucleus profiling of the P4 hypothalamus reveals sex-resolved and cell type-specific pathway remodeling in *Magel2* KO mice. **(A)** Schematic overview of the computational workflow. Hypothalami were isolated from postnatal day 4 (P4) WT and *Magel2* KO (both sexes) and processed for single-nucleus RNA sequencing (snRNA-seq). Following preprocessing, integration, clustering, and cell-type annotation, downstream analyses included differential expression, functional enrichment, microglial state classification, and cell-cell communication inference. **(B-C)** UMAP visualization of integrated datasets colored by cell type for female (B) and male (C) samples, showing consistent identification of major hypothalamic populations, including excitatory and inhibitory neurons, astrocytes, oligodendrocyte lineage cells, ependymal cells, endothelial cells, cycling cells, and microglia. **(D-E)** Cell type-resolved GSEA heatmaps for female (D) and male (E) samples summarizing pathway-level alterations between *Magel2* KO and WT across excitatory neurons, inhibitory neurons, astrocytes, OPCs, and microglia. Columns indicate the cell types in which normalized enrichment score (NES) values are displayed, whereas rows represent representative pathways selected from the GSEA results of the cell type indicated by the row annotation. For each selected pathway, NES values are shown across all analyzed cell types to distinguish cell type-restricted from shared transcriptional programs. Positive NES values indicate enrichment in *Magel2* KO relative to WT, whereas negative NES values indicate enrichment in WT relative to *Magel2* KO. Asterisks denote significantly enriched pathways (adjusted *q* < 0.05). Red pathway labels indicate pathways that were significantly altered in more than one cell type, highlighting recurrent programs shared across populations. Black boxes outline the matched cell type blocks from which representative pathways were selected.

To investigate whether *Magel2* deletion affects transcriptional pathways across different hypothalamic cell types, cell type-resolved GSEA analysis was performed separately in female and male datasets. This analysis revealed pathway-level alterations across several cell populations, including shared changes in translation- and ribosome-associated programs (**Figure 1D, E**). However, microglia showed a particularly strong pattern of significant pathway alteration in both sexes, as indicated by the number of significantly enriched terms in the microglial column. In females, microglial changes were mainly associated with synapse-related pathways, including presynaptic and postsynaptic membrane-associated terms (**Figure 1D**). In males, altered microglial pathways included ion channel activity, glutamate receptor activity, and regulation of presynaptic membrane potential (**Figure 1E**). Based on these findings, we focused our subsequent analyses predominantly on microglia, as snRNA-seq identified them as the most profoundly transcriptionally dysregulated cell population in the *Magel2* KO hypothalamus.

Given the close relationship of microglial morphology and function [23–25], we aimed to determine whether *Magel2* deficiency alters the structural characteristics of microglia. We first performed high-resolution three-dimensional morphometric analysis of hypothalamic microglia in *Magel2* KO and WT mice [26–27] (**Supplementary Figure 2A, B)**, which enables detailed three-dimensional reconstruction of microglial surfaces and processes. Structural differences were readily apparent, with microglia from *Magel2* KO mice displaying both altered surface morphology and reduced process complexity (**Figure 2A**). These observations were quantitatively supported by significant reductions in surface area, process length, and peak Sholl intersection values (**Figures 2B-D**). Conventional Sholl analysis further revealed that microglia in *Magel2* KO mice exhibit reduced spatial domain (i.e. surveillance territory) and markedly decreased arborization complexity (**Figure 2E**). To complement these findings, we generated multidimensional heatmaps that integrate domain size and Sholl intersection visualizing both spatial extent and progressive branching complexity (**Figure 2F, G**). Using this approach, we confirmed that *Magel2* KO microglia occupy a smaller three-dimensional territory and are substantially less complex (**Figure 2G**).

**Figure 2.**
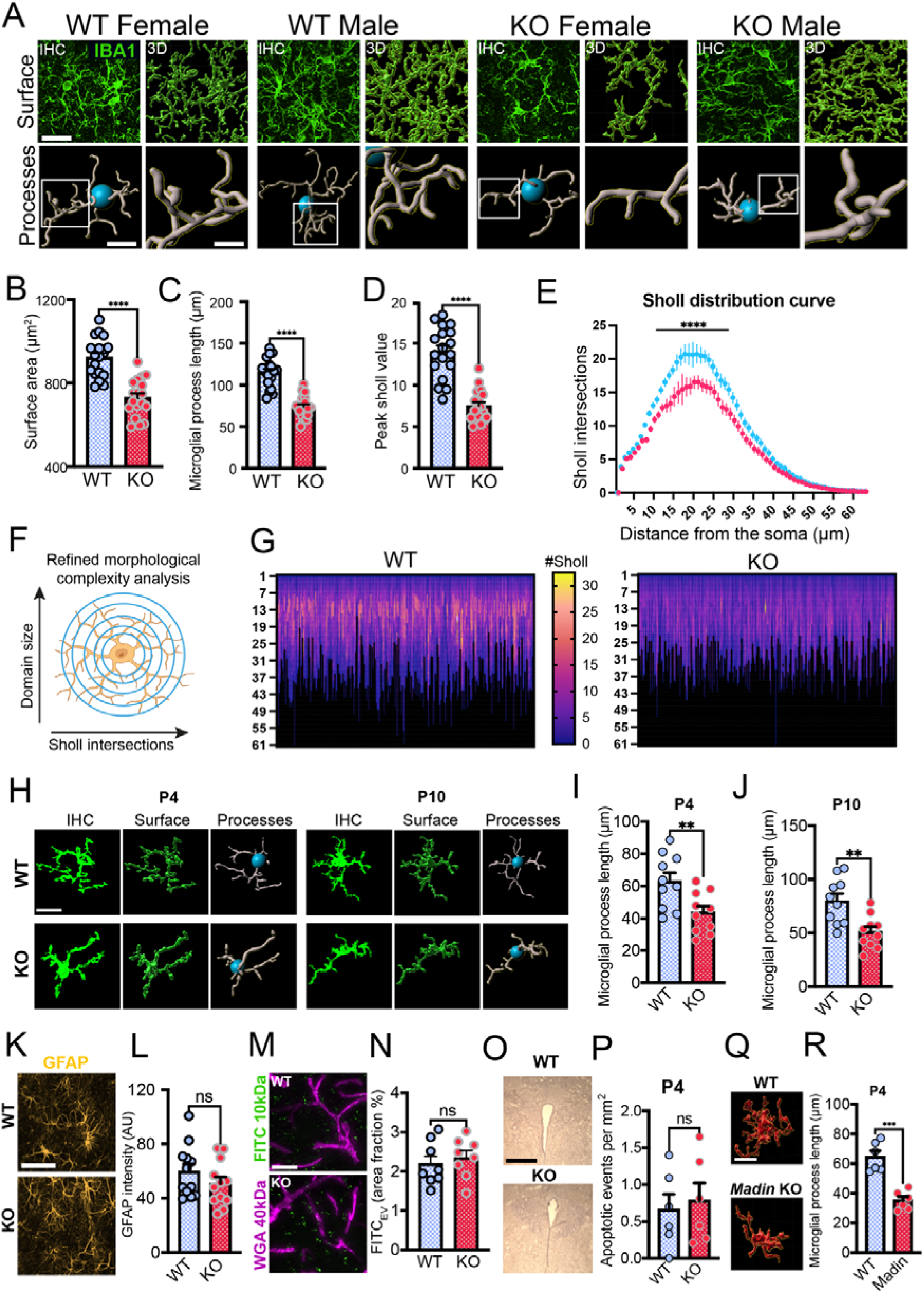
Alterations of microglial morphology in *Magel2* KO mice. **A** Overview images highlight microglial distribution within the hypothalamus of WT and *Magel2* KO mice. Enlarged insets show altered microglial morphology based on staining against the microglial marker IBA1. Following three-dimensional reconstruction, microglia from Magel2 KO mice display visibly reduced microglial process complexity. Scale bars = 200 µm, 20 µm and 5 µm. **B** Reduced microglial surface area in *Magel2* KO mice, n=18 per group (9 male / 9 female). **C** Reduced microglial process length in *Magel2* KO mice, n=18 per group (9 male / 9 female). **D** Peak sholl value reduction in microglia from *Magel2* KO mice. **E** *Magel2* KO mice display significant alterations in the microglial sholl distribution curve. Two-way ANOVA, posthoc Tukey test, ****p<0.0001. **F** Principle of Sholl intersection for the three-dimensional assessment of microglial morphology. Intersections of artificially introduced spheres with microglial processes are quantified, the peak sholl value refers to the maximum complexity of a given microglial cell. **G** Heatmaps highlight microglial domain size and peak sholl values of 256 randomly sampled microglia from WT and *Magel2* KO mice. **H** Representative confocal images depicting microglia from WT and *Magel2* KO mice at P4 and P10. Scale bar = 10 µm. **I** Reduced microglial process length in P4 *Magel2* KO mice. **J** Reduced microglial process length in P10 *Magel2* KO mice. **K, L** Immunohistochemical assessment of GFAP signal reactivity in WT and Magel2 KO mice, n=12 per group (6 male / 6 female). Scale bar = 100 µm. **M, N** Labeling of brain vasculature and assessment of BBB integrity via perfusion with FITC 10 kDa and WGA 40 kDa. Graph shows extravasated FITC_EV_ quantified via area fraction (%) in WT and *Magel2* KO mice, scale bar = 100 µm. **O, P** Quantification of apoptotic events in the hypothalamus of WT and *Magel2* KO mice at P4, n=6 per group (3 male / 3 female). **Q** Representative examples of typical microglia in Madin and WT mice at P4. **R** Microglia from Madin mice display significantly reduced process length at P4, n=6 per group (3 male / 3 female). Scale bar = 10 µm. Unpaired, two-tailed t-test, ****p<0.0001. Unpaired, two-tailed t-test, **p<0.01, ****p<0.0001.

In parallel, we assessed physiological parameters and found that *Magel2* KO mice exhibited significantly increased body weight, abdominal circumference and enlargement of gonadal fat pads compared to WT controls (**Supplementary Figure 2C-E**). In *Magel2* KO mice - but not in WT - a robust and statistically significant negative correlation was detected between body weight and microglial complexity (p < 0.0001, R² = 0.6219, **Supplementary Figure 2F, G**). To determine whether the observed microglial morphological changes were influenced by increased body weight, we analyzed microglia at earlier developmental stages (postnatal day 4 and day 10), when *Magel2* KO mice are underweight and developmentally delayed relative to WT [28]. We observed similar trends toward reduced microglial complexity at early time points (P4 and P10) (**Figure 2H-J**), suggesting that the loss of microglial complexity is not merely a consequence of increased body mass, but rather reflects an intrinsic microglial alteration associated with *Magel2* deficiency.

Since the microglial deramification observed in *Magel2* KO mice resembles changes seen during sustained neuroinflammation, we next asked whether hallmarks of a classical inflammatory cascade, including blood-brain barrier disruption [29], astrocyte reactivity [30], and apoptotic cell death, were present. We performed immunohistochemical analyses to examine potential pathological changes in astrocyte reactivity (**Figure 2K, L**), but detected no difference between the two groups. To probe for potential BBB compromise, we conducted transcardiac perfusions using FITC-dextran (10 kDa) and WGA-dextran (40 kDa), dyes that respectively extravasate into the brain parenchyma or selectively label the cerebral vasculature. We observed comparable FITC-dextran extravasation between groups (**Figure 2M, N**), suggesting that the BBB integrity is preserved in *Magel2* KO mice. Next, we quantified the number of apoptotic events within the hypothalamus at the developmental apoptosis peak P4, but found no differences between the groups (**Figure 2O, P**). *Magel2* and *Necdin* are co-expressed and co-regulated within the PWS locus, with deletion of one gene altering the expression of the other, suggesting functional interplay between these imprinted genes [31]. To test whether alterations in microglial morphology are a unique feature of *Magel2* KO mice, we utilized the recently generated *Madin* mouse, which lacks both *Magel2* and *Necdin,* and assessed microglia at P4 (**Figure 2Q, R**). We observed a significant reduction in process length that appeared to be even more pronounced than in the *Magel2* KO mice (**Figure 2J**). To further probe whether alterations in microglia morphology were a regional or brain-wide phenomenon, we assessed the nucleus accumbens, dorsal CA2 of the hippocampus and the caudate putamen as anatomically and functionally distinct, non-hypothalamic regions not primarily implicated in PWS pathology. In neither of these brain regions, surface area or process length was significantly different between WT and *Magel2* KO mice (**Supplementary Figure 3A-F**), suggesting that the microglial morphological alterations may be confined to the hypothalamus in this model.

Based on the findings, we hypothesized that microglia in *Magel2* KO mice may be altered in their function as was previously indicated by analysis of a human bulk transcriptomic dataset [11]. Differential expression analysis revealed substantial transcriptional remodeling in microglia following *Magel2* deletion (**Figure 3A, B**). In these microglia, numerous genes involved in immune signaling, cellular communication, and regulatory processes were significantly altered relative to WT controls, with the magnitude of these changes varying by sex. One noteworthy observation was the consistent upregulation of Xist across cell types in female mice. To validate these findings, we performed Xist RNA-FISH [32] on hypothalamic samples from WT and *Magel2* KO mice at P4 but did not detect differences in Xist signal intensity (**Supplementary Figure 4A, B**). Notably, substantial variability in Xist signal was observed both between animals and across individual cells (**Supplementary Figure 4C**), which may account for the discrepancy between snRNA-seq and RNA-FISH results. Of note, we observed substantial and consistent transcriptomic changes across all analyzed cell types in Magel2 KO mice (**Supplementary Figures 5-8**).

**Figure 3.**
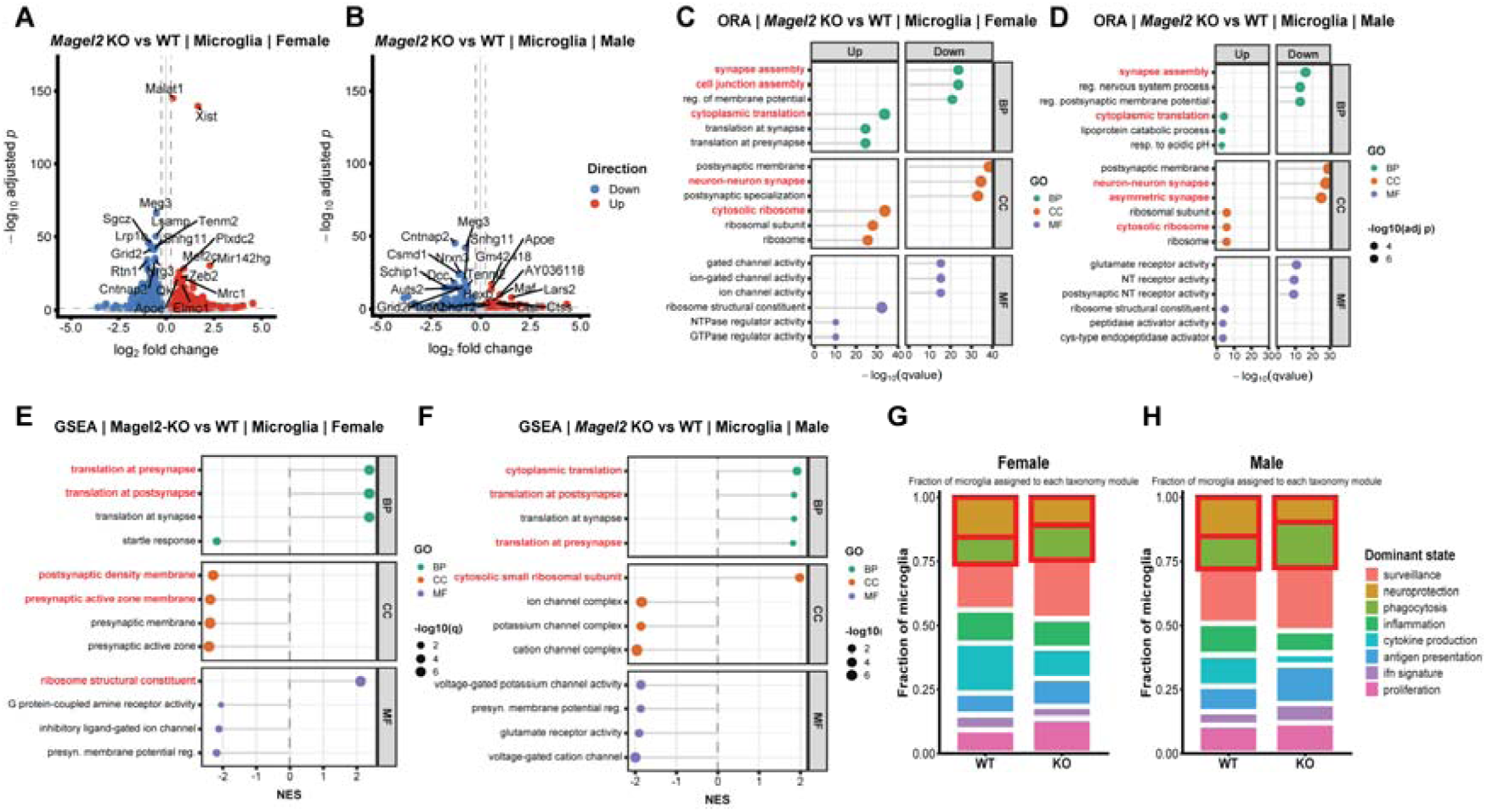
*Magel2* deletion remodels the microglial transcriptome toward phagocytic, inflammatory, translational, and synapse-associated programs. **(A-B)** Volcano plots of differentially gene expression between *Magel2* KO and WT microglia in female(A) and male(B) datasets. Differential expression was computed using pseudo-bulk approach based on aggregated counts and fractional expression. Statistical significance was assessed using Fisher‘s exact test followed by Benjamini-Hochberg (BH) correction. Genes with absolute log2FC ≥0.25 and adjusted p (padj) <0.05 are highlighted (red: upregulated in KO; blue: downregulated in KO). **(C-D)** Gene Ontology over-representation analysis (ORA) of significantly upregulated and downregulated genes in microglia from female (C) and male(D). Dot size represents gene ratio, and color indicates adjusted p-value significance. Enriched pathways highlight immune activation, inflammation signaling, lysosomal and phagocytosis-related processes, and alterations in cellular homeostasis **(E-F)** Gene set enrichment analysis (GSEA) of ranked microglial transcriptional changes in female(E) and male(F) datasets. Normalized enrichment scores (NES) are shown for representative GO terms across biological process, cellular components, and molecular function categories, revealing coordinated upregulation of immune and translational programs alongside downregulation synapse-associated and neuronal signaling pathways. **(G-H)** Microglial taxonomy analysis in female(G) and male(H) samples based on module scoring of predefined microglial state signatures. Stacked bar plots show the fraction of microglia assigned to each dominant state, including surveillance, neuroprotection, phagocytosis, inflammation, cytokine production, antigen presentation, interferon response, proliferation. *Magel2* KO samples exhibit a shift toward phagocytotic and inflammatory states with a relative reduction of homeostatic/surveillance-associated programs.

Functional enrichment analyses further indicated that *Magel2* loss affects biological pathways associated with microglial activation and intercellular signaling, processes that are critical for microglia-neuron communication. Over-representation analysis and gene set enrichment analysis converged on a consistent pattern of transcriptional reprogramming in *Magel2* KO microglia (**Figure 3C-F**). Both approaches revealed enrichment of pathways related to regulation of synaptic activity, trans-synaptic signaling, and membrane potential, together with cellular component terms centered on synaptic structures such as neuron-to-neuron synapse, postsynaptic density, and presynaptic membrane, as well as molecular functions including ion channel activity and neurotransmitter receptor activity. In parallel, GSEA demonstrated positive enrichment of cytoplasmic translation and ribosome-associated processes, indicating increased biosynthetic and metabolic activity. Conversely, both ORA (from downregulated genes) and GSEA consistently highlighted a depletion of pathways associated with synaptic organization, ion channel complexes, and neuronal signaling, including presynaptic and postsynaptic membrane components and voltage-gated ion channel activity. Together, these results define a coherent shift in *Magel2* KO microglia toward a state characterized by enhanced translational capacity and altered immune-associated signaling, accompanied by a relative loss of transcriptional programs linked to synaptic interaction and neuronal communication, suggesting a functional decoupling of microglia from neuronal networks [33–35].

Interestingly, the magnitude of transcriptional changes differed between sexes, consistent with previous reports [36]. Female *Magel2* KO microglia exhibited a larger number of differentially expressed genes compared with male *Magel2* KO microglia, suggesting that *Magel2*-dependent regulation of microglial transcriptional programs may be more pronounced in females. To place these transcriptional changes into a broader functional framework, we next classified microglia according to recently defined taxonomy modules derived from cross-condition transcriptomic microglia states (**Figure 3G, H**, [37]). This analysis showed that *Magel2* loss did not collapse microglia into a single dominant pathological identity, but instead redistributed cells across multiple functional modules. In both sexes, *Magel2* KO samples showed a relative reduction in the neuroprotection-associated module and a concomitant redistribution toward surveillance-, cytokine production-, interferon signature-, and proliferation-associated states, with additional shifts in phagocytosis- and inflammation-related modules that differed in magnitude between female and males. Thus, the taxonomy analysis supports the conclusion that *Magel2* loss perturbs the balance of homeostatic versus reactive microglial states rather than producing a uniform activation profile. Given that these modules were defined from a large cross-species and cross-pathology transcriptomics framework, their redistribution in the *Magel2* KO hypothalamus further suggests that this model engages biologically meaningful axes of microglial state plasticity [37]. Next, we compared our snRNA-seq data with a previously published bulk transcriptomic dataset from PWS individuals [11] and observed remarkable overlap in numerous genes and pathways, most notably the upregulation of complement cascade genes (i.e C1qa, C1qb and C1qc, **Supplementary Figure S9A-E**) as well as the absence of transcriptional signatures associated with neuroinflammation and apoptosis **(Supplementary Figure S9F, G**).

Microglia form specialized contacts with neuronal somata that enable direct monitoring and modulation of neuronal function [17–19]. To specifically investigate potential alterations in microglia-neuron communication, we performed co-staining for IBA1 and the neuronal marker NeuN (**Figure 4A**) as a proxy for microglia-neuron communication to avoid confounding contacts with neuronal membranes and dendrites in our subsequent Imaris-based analysis. Using our established Imaris-based pipeline, we performed three-dimensional reconstruction of microglial and neuronal surfaces. We first quantified microglial and neuronal cell numbers in the hypothalamus of WT and *Magel2* KO mice to determine whether normalization of interactions was necessary (**Figure 4B, C**). Average counts of microglia and neurons did not differ significantly between groups (**Figure 4B, C**). Next, we co-stained brain sections for IBA1 and neuronal marker TUBB3 to investigate the type of microglia-neuron interactions (**Figure 4D**). Two distinct types of contacts were identified: *appositional*, involving elongated microglial processes spanning neuronal membranes, and *focal*, characterized by concentrated contacts with slightly enlarged interfaces. Notably, the frequency of focal microglial contacts with neuronal membranes was elevated 2.2-fold in *Magel2* KO mice (**Figure 4E**). We further observed that the proportion of neuronal somata in contact with microglial processes (defined as microglial interactions) was unaltered between the groups. Interestingly, despite the previously reported loss of microglial complexity (**Figure 1**) and the fact that the percentage of neurons in contact with microglial processes was unaltered (**Figure 4F**), we observed an increase in microglia-neuron contact intensity in *Magel2* KO mice, as measured by surface area overlap (**Figure 4G**). We hypothesized that *Magel2* KO microglia may be in a chronic state of activation resulting in altered communication with neurons. Following sustained neuroinflammation, microglia may engage in CD68-dependent phagocytosis of various components of the BBB including astrocytic endfeet [29]. While we did not observe alterations in astrocytic coverage or BBB integrity (**Figure 1**), we observed a trend towards increased CD68 expression in microglia from *Magel2* KO mice (**Figure 4H-J**). Microglia play a critical role during early brain development by dynamically interacting with synaptic elements to regulate synaptic pruning and remodeling, thereby shaping neuronal circuit maturation [20, 33, 35]. Thus, we speculated that microglia in *Magel2* KO mice may be involved in excessive synaptic pruning and co-stained brain sections with postsynaptic marker PSD95 at P14 (**Figure 4K**), where developmental pruning is at its peak [33, 35]. Intriguingly, we observed a reduction of PSD95 density [38] (**Figure 4L**) accompanied by an increase of engulfed PSD95 [39] in microglia in *Magel2* KO mice (**Figure 4M**).

**Figure 4.**
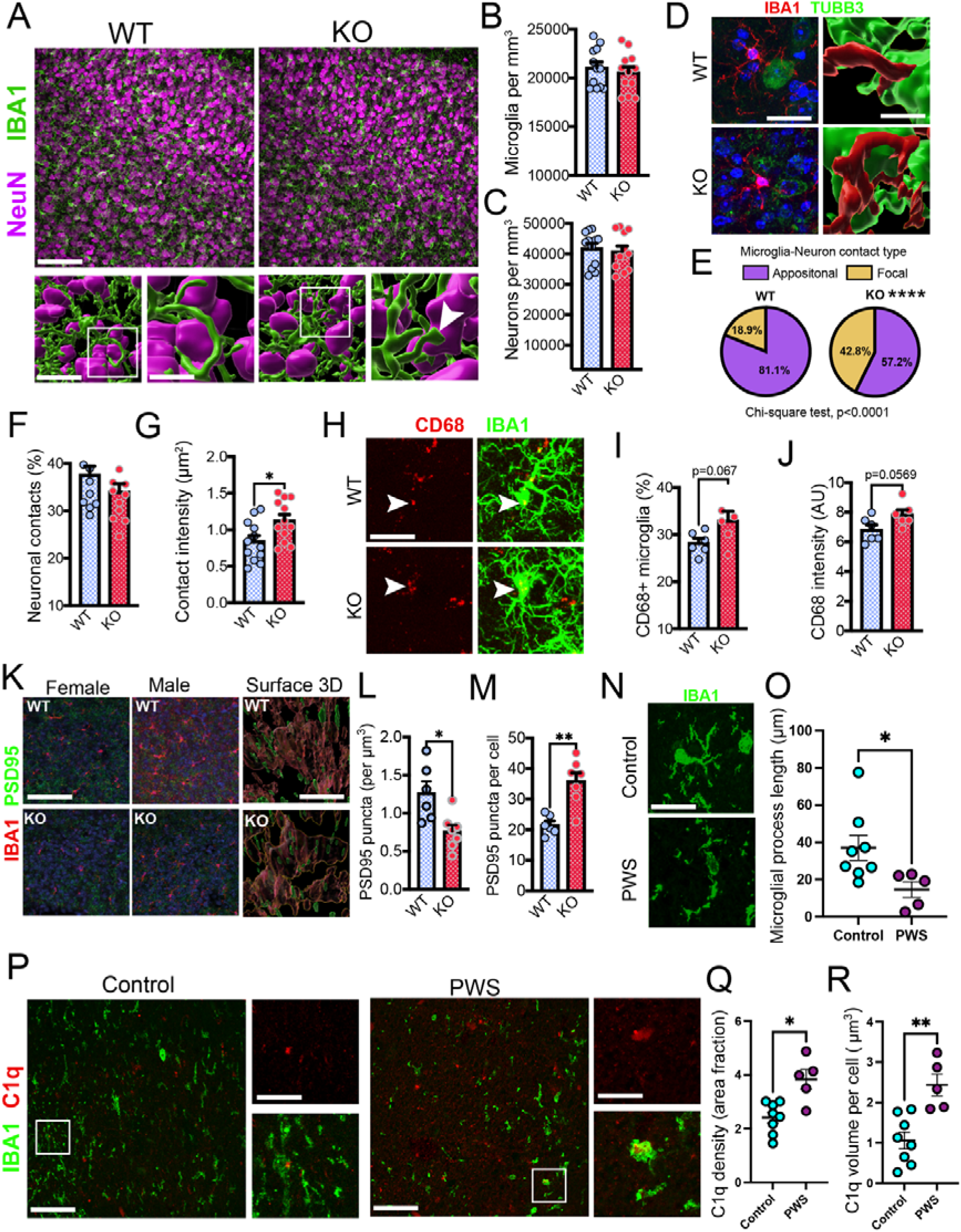
Altered microglia-neuron contacts and increased synaptic pruning in *Magel2* KO and post-mortem PWS brain samples. **A** Co-staining for IBA1 and NeuN in the hypothalamus of WT and *Magel2* KO mice. High magnification insets show intensified microglia-neuron contacts in *Magel2* KO mice. Scale bars = 100 µm, 5 µm and 2 µm. **B** Assessment of microglia density in WT and *Magel2* KO mice, n=12 per group (6 male / 6 female). **C** Assessment of neuronal density in WT and *Magel2* KO mice, n=12 per group (6 male / 6 female). **D** Co-staining for IBA1 and TUBB3 in the hypothalamus of WT and *Magel2* KO mice. Scale bar = 25 µm. Three-dimensional reconstruction highlights intensified microglia-neuron interactions in *Magel2* KO mice. Scale bar = 1 µm. **E** Pie charts quantify the relative contributions of appositional and focal contacts in WT and *Magel2* KO mice, n=12 per group (6 male / 6 female). Chi-square test, p<0.0001. **F** Quantification of neuronal contacts in WT and *Magel2* KO mice, defined as neurons in contact with microglial processes, n=12 per group (6 male / 6 female). **G** Contact intensity of microglia-neuron contacts, defined as overlapping surface of IBA1 and TUBB3 signals, n=12 per group (6 male / 6 female). **H** Co-staining against IBA1 and CD68. White arrowheads indicate co-expression of CD68 and IBA1 signals. Scale bar = 25 µm. **I** Percentage of CD68-positive microglia in WT and *Magel2* KO mice, n=6 per group (3 male / 3 female). **J** Relative CD68 signal intensity within microglia of *Magel2* KO and WT mice measured in AU, n=6 per group (3 male / 3 female). **K** Immunohistochemical staining against IBA1 and PSD95. Representative images highlight increased PSD95 puncta in the hypothalamus of male and female *Magel2* KO mice. Three-dimensional reconstruction shows PSD95 engulfment by IBA1-positive microglia (made transparent for visual purposes). Note the increase in engulfed PSD95-positive puncta in the *Magel2* KO microglia. Scale bars = 100 µm and 5 µm. **L** Quantification of PSD95 puncta per mm^3^, n=6/7 per group (WT: 3 male / 3 female, KO: 3 male / 4 female). **M** Quantification of PSD95 puncta within microglia of *Magel2* KO and WT mice n=6/7 per group (WT: 3 male / 3 female, KO: 3 male / 4 female). **N** IBA1 staining in post-mortem hypothalamic brain samples of control subjects and PWS Type 2 individuals. Scale bar = 10 µm. **O** Microglia from PWS Type 2 samples show significantly reduced process length, n=8 control subjects and n=5 PWS subjects. **P** Immunohistochemical staining against IBA1 and C1q in hypothalamic post-mortem brain samples. High magnification insets show C1q co-expression in IBA1-positive microglia. Scale bars = 100 µm and 10 µm. **Q** Quantification of C1q signal intensity in hypothalamic post-mortem samples, n=8 control subjects and n=5 PWS subjects. **R** Quantification of microglia-bound C1q levels (per cell), n=8 control subjects and n=5 PWS subjects. Unpaired, two-tailed t-test, *p<0.05, **p<0.01.

To probe the translational relevance of our findings obtained in mice, we processed post-mortem hypothalamic brain samples from PWS individuals and age-matched controls. We observed that microglia from PWS individuals often appeared fragmented, with short and disorganized processes (**Figure 4N**). When we applied our three-dimensional reconstruction, we observed reduced process length (**Figure 4O**) consistent with previous reports [12]. Next, given that the human hypothalamic synaptic landscape is largely unknown, we co-stained brain sections with IBA1 and C1q (**Figure 4P**), as the complement cascade critically involved in synaptic pruning [40] and analyzed C1q signals in analogy with previous studies [41–43]. We observed significantly increased C1q levels, along with drastically elevated microglia-bound C1q in microglia from PWS individuals (**Figure 4Q, R**). Together, increased C1q engagement in human PWS microglia and enhanced PSD95 engulfment in *Magel2* KO mice indicate a shift toward a phagocytic microglial state, consistent with the transcriptional reprogramming identified by snRNA-seq (**Figure 3**).

Based on these observations, we next investigated whether *Magel2* deletion alters intercellular communication within the brain microenvironment by applying CellChat to infer ligand-receptor-mediated signaling networks from our mouse snRNA-seq transcriptomic data [44]. This approach enabled a systematic comparison of communication patterns between WT and *Magel2* KO conditions, with a focus on microglia interactions with neuronal and glial populations. Global network analysis revealed genotype-dependent remodeling of cell-cell communication across brain cell types (**Figure 5A, B**). In females, *Magel2* deletion was associated with a broader and denser predicted communication network, reflected by an increased number of inferred interactions across multiple cell populations (**Figure 5A**). This increase was particularly prominent in interactions involving neurons and oligodendroglial lineage cells, suggesting a broad reorganization of the neuro-glial communication landscape [45]. This network-level expansion was particularly evident for microglia, which showed a marked increase in both incoming and outgoing interaction numbers in *Magel2* KO compared with WT samples (**Figure 5E**). In males, genotype-dependent remodeling was also detected, but the pattern appeared more restricted: microglia showed a clear increase in incoming interactions, whereas outgoing microglial interactions remained largely unchanged between WT and KO conditions (**Figure 5B, F**). Thus, *Magel2* loss remodels intercellular communication in both sexes, but the female dataset shows a more broadly expanded microglial communication phenotype. At the pathway level, RankNet analysis identified specific signaling programs driving this network remodeling (**Figure 5C, D**). In female *Magel2* KO sample, the most prominently altered pathways were dominated by cell adhesion and synaptic interactions systems, including NCAM, CADM, CDH, CNTN, NRXN, NGL, and NEGR signaling. These pathways are key mediators of direct cell-cell contact, particularly at synaptic and peri-synaptic interfaces, and are critical for maintaining neuronal connectivity and neuron-glia interactions [18–19, 35]. In addition, axon guidance-related pathways, such as SEMA6 and Ephrin signaling (EPHA/EPHB), were also differentially regulated, suggesting alterations in structural communication cues that influence circuit organization. Alongside these contact-dependent systems, NRG (neuregulin) and PTN (pleiotrophin) signaling pathways were enriched, indicating changes in trophic communication between neurons and glial cells. Together, these results indicate that *Magel2* loss primarily affects contact-mediated and synapse-associated communication pathways, rather than extracellular matrix-driven signaling, pointing to a reorganization of intercellular signaling at the level of neuronal connectivity and microglial interaction interfaces.

**Figure 5.**
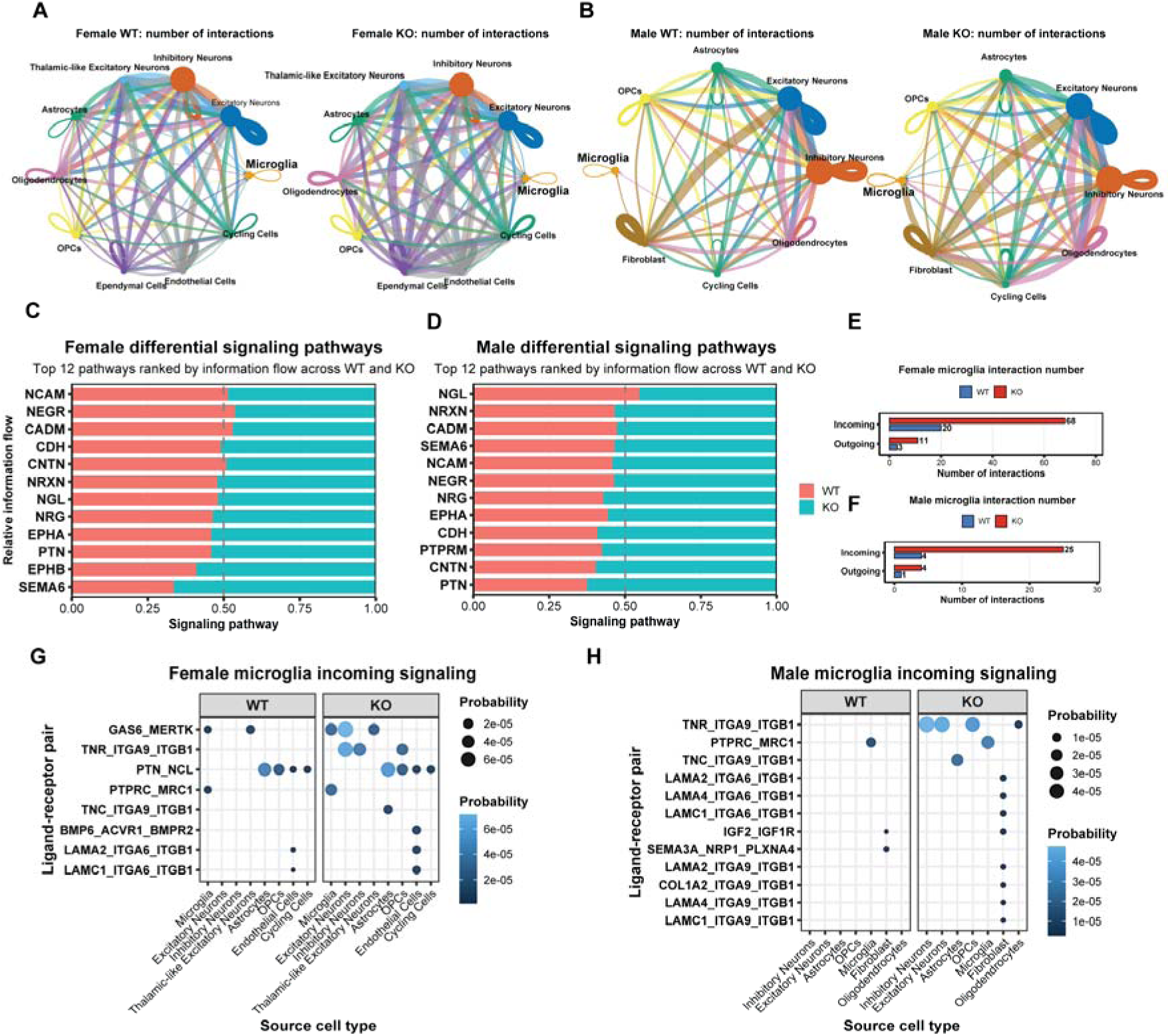
*Magel2* deletion remodels intercellular communication and enhances microglia-directed signaling in the P4 hypothalamus. **(A-B)** CellChat-based circle plots showing the predicted number of ligand–receptor-mediated interactions across hypothalamic cell types in female (A) and male (B) WT and *Magel2* KO datasets. Edge thickness represents the number of predicted interactions between cell populations, and node size reflects the relative abundance of each cell type. Global communication networks show genotype-dependent changes in the overall interaction structure across neural and glial populations. **(C-D)** Differential signaling pathway analysis in female (C) and male (D) datasets. The top 12 signaling pathways were ranked based on relative information flow across WT and *Magel2* KO conditions. Stacked bars indicate the relative contribution of each genotype to the total information flow for each pathway, highlighting genotype-associated remodeling of major communication pathways, including NCAM, NEGR, CADM, CNTN, NRXN/NGL, NRG, EPHA, PTN, and SEMA-related signaling. **(E-F)** Quantification of microglia-associated interaction numbers in female (E) and male (F) datasets. Incoming interactions indicate signaling directed toward microglia, whereas outgoing interactions indicate signaling from microglia to other cell types. *Magel2* KO samples show increased microglia-directed incoming interactions in both sexes, with a stronger effect observed in females. **(G-H)** Bubble plots showing microglia incoming ligand–receptor interactions in female (G) and male (H) datasets. Columns indicate source cell types, rows indicate ligand-receptor pairs, and WT and *Magel2* KO conditions are shown separately. Dot sizes represent the inferred CellChat communication probability for each ligand-receptor pair. *Magel2* KO microglia receive increased predicted signaling from multiple source populations, including neuronal, astrocytic, OPC, endothelial, fibroblast, and oligodendrocyte-lineage populations, with prominent changes in extracellular matrix-, adhesion-, trophic-, and immune-related ligand-receptor interactions. Cell-cell communication analysis was performed using CellChat separately for WT and *Magel2* KO datasets in each sex.

Microglia-centered analyses further demonstrated that *Magel2* deficiency reshapes signaling interactions between microglia and surrounding cell populations (**Figure 5E, F**). This is indicative of enhanced environmental sensing rather than active signal broadcasting. In line with this observation, incoming signaling analysis (**Figure 5G**) revealed that microglia receive a distinct set of ligand-receptor interactions enriched for integrin-mediated adhesion, trophic signaling, and immune-related pathways. In females, incoming signals prominently included PTN-NCL, GAS6-MERTK, and TNF- and TNC-associated integrin interactions, indicating enhanced trophic support, phagocytic signaling, and adhesion-mediated communication. Additional contributions from LAMININ-integrin interactions were also observed, although these were less dominant compared to trophic and immune-associated signals [46]. In males, microglial incoming signaling was more strongly dominated by extracellular matrix-associated interactions, particularly multiple LAMININ-integrin and COLLAGEN-integrin (COL1A2) ligand-receptor pairs, suggesting a greater reliance on structural and matrix-mediated signaling inputs. In addition, SEMA3A-NRP1-PLXNA4 and IGF2-IGF1R interactions indicate concurrent engagement of axon guidance and growth factor signaling pathways.

Together, these data demonstrate that *Magel2* plays a critical role in regulating cell-cell interaction and ligand-receptor-mediated communication within the hypothalamus. Its loss leads to a reorganization of intercellular signaling networks characterized by increased incoming signaling to microglia, with distinct qualitative differences between sexes. In females, this shift is primarily driven by enhanced trophic and immune-associated signaling, including pathways linked to phagocytic regulation and microglial activation. In contrast, male microglia exhibit a stronger enrichment of integrin-mediated extracellular matrix interactions, indicating a greater reliance on structural and adhesion-related signaling inputs [46]. Collectively, these findings suggest that *Magel2* deficiency promotes a state of increased microglial signal reception, consistent with altered responsiveness to environmental cues. This reorganization of communication networks is likely to impact how microglia integrate neuronal and glial signals, with potential consequences for synaptic regulation and neuroimmune interactions in a sex-dependent manner.

## Discussion

Here, we show that *Magel2* KO mice display pronounced loss of microglial complexity, increased focal microglia-neuron contacts, and elevated complement engagement, supported by 3D Imaris-based reconstructions and human post-mortem tissue validation. Microglial abnormalities observed in our *Magel2* KO model are consistent with growing evidence that neuroinflammation contributes to PWS pathophysiology [10–12]. Dysregulated microglial morphology and elevated microglia-neuron interactions in *Magel2* KO mice may impair hypothalamic neuronal development, synaptic organization and function, especially during early developmental stages, which could explain, at least partially, PWS neurocircuitry alterations. Based on our data, it seems plausible that excessive weight gain further exacerbates microglial deramification, although morphological alterations already appeared at P4 and P10, when no weight changes were observed, highlighting a weight-independent effect.

Neuroinflammatory signatures have been reported in PWS [11], including elevated circulating cytokines and hypothalamic transcriptional dysregulation [47]. Interestingly, BMI-matched controls displayed significantly lower circulating cytokines compared with PWS patients [47], suggesting that these neuroinflammatory alterations are at least partly weight-independent. Our data support that microglial morphological and molecular abnormalities are intrinsic to *Magel2* deficiency, rather than a consequence of the obesity phenotype in PWS. Our findings also align with observations that *Magel2* deficiency disrupts hypothalamic homeostasis [48] and may promote maladaptive microglial activation. Together, this supports a model in which aberrant microglial function contributes to synaptic and metabolic phenotypes in PWS and suggests *Magel2* deficiency drives an early shift in microglial state characterized by reduced structural complexity and enhanced engagement with neuronal elements, which may alter synaptic remodeling during hypothalamic circuit development.

A previous study identified microglial morphological and transcriptional differences between Type 1 and 2 PWS deletions and suggested that *Cyfip1*, a gene only affected by the larger deletion (BP1), may partly account for these differences [12]. Interestingly, we see substantial microglial alterations in our PWS Type 2 cohort, suggesting that effects may even be more pronounced in individuals with PWS Type 1. Our study demonstrates early-life microglial deramification independent of weight gain and provides the first evidence of increased C1q engagement in human PWS hypothalamus. Collectively, our data support a model in which *Magel2* deficiency drives a shift toward a phagocytic microglial state characterized by increased complement engagement and synaptic engulfment, consistent with established roles of C1q-dependent pathways in activity-dependent synaptic pruning [40–42]. These findings raise exciting therapeutic possibilities: microglial dampening strategies such as minocycline, which reduces microglial activation and has shown benefit in mouse neuroinflammation models [49], may be used to ameliorate microglia-associated neuroinflammation in PWS patients. Additionally, CSF1R inhibitors like PLX5622 have been shown to modulate microglial number and state in mice without causing major behavioral deficits, however they are not microglia specific and can impact macrophage cells [50]. Based on our data, targeting the complement cascade [51] directly may be more mechanistically precise than broadly inhibiting microglia, as C1q/C3-dependent synaptic pruning drives engulfment and its blockade rescues synapse loss [40, 42]. In contrast, nonspecific microglial inhibition may impair homeostatic functions without effectively blocking complement-mediated pruning.

Our snRNA sequencing analysis revealed transcriptional changes in microglia that are consistent with altered synaptic organization and pruning processes [33, 35, 43, 52]. In particular, enrichment of pathways related to immune signaling and complement activation suggests a shift toward a more phagocytic, synapse-engulfing phenotype, aligning with the observed increase in C1q engagement and PSD95 uptake. Together, these data provide direct structural and transcriptional evidence that *Magel2* deficiency enhances microglia-mediated synaptic engulfment, linking transcriptional changes to functional alterations in synaptic remodeling. In parallel, differential expression of genes involved in cellular communication and environmental sensing indicates that microglia may respond differently to neuronal cues, potentially disrupting activity-dependent synaptic refinement.

Consistent with this interpretation, our CellChat analysis revealed a substantial reorganization of ligand-receptor-mediated communication networks, particularly affecting pathways involved in cell-cell contact and signal reception. At the systems level, signaling changes were dominated by adhesion- and synapse-associated pathways, including NCAM, NRXN, CADM, and CNTN families, which are critical for maintaining synaptic structure and neuron-glia interactions [53–54]. These findings suggest that *Magel2* deficiency alters the molecular interface through which microglia interact with neuronal elements, potentially modifying how synaptic signals are recognized and processed. At the level of microglial signal integration, incoming communication analysis further demonstrated that microglia in *Magel2* KO hypothalamus receive enhanced inputs through trophic, immune, and adhesion-related signaling pathways. In particular, enrichment of PTN-mediated signaling, GAS6-MERTK interactions, and integrin-dependent adhesion cues that microglia are exposed to may favor a microenvironment that promotes both structural engagement and functional activation. MERTK signaling is a well-established regulator of microglial phagocytosis and synaptic engulfment [55–56] while PTN and related growth factor pathways have been implicated in neuron-glia communication and synaptic plasticity. Integrin-mediated signaling further contributes to microglial adhesion, migration, and interaction with synaptic structures [57–58]. Together, these signaling inputs provide a mechanistic framework linking altered intercellular communication to the enhanced synaptic pruning phenotype observed in *Magel2*-deficient hypothalamus. Interestingly, the nature of incoming signals differed between sexes. In females, microglial signaling was more strongly associated with trophic and immune-related pathways, suggesting enhanced responsiveness to neuronal and damage-associated cues. In contrast, male microglia showed a relative enrichment of integrin-mediated extracellular matrix interactions, indicating a shift toward structural and adhesion-driven signaling. These findings are consistent with emerging evidence that microglial states and signaling responsiveness are strongly influenced by sex-specific developmental trajectories and hormonal regulation [59–60]. Thus, *Magel2* loss may engage distinct microglial activation programs depending on sex, potentially influencing how microglia interpret and respond to their microenvironment.

The enrichment of pathways linked to ribosomal function and protein translation points to broader alterations in microglial cellular homeostasis, which could impair their capacity to properly regulate synaptic remodeling. Notably, the more pronounced transcriptional and communication changes observed in females suggest that *Magel2*-dependent regulation of microglial function may be influenced by sex-specific developmental or hormonal factors. Given established sex differences in microglial maturation and activation states [36], these findings point to a potential sex-dependent component of microglial dysregulation in PWS. Together, these transcriptional signatures support a model in which *Magel2* deficiency drives microglia toward a state that may promote excessive or dysregulated synaptic pruning, thereby contributing to impaired neural circuit development. The long non-coding RNA *Xist* plays a crucial role in the epigenetic process of X chromosome inactivation [61]. It remains unclear why differential Xist expression was detected across all hypothalamic cell populations in females by snRNA-seq but could not be validated by RNA-FISH. The discrepancy between snRNA-seq and RNA-FISH likely reflects methodological differences and substantial inter-cellular and inter-individual variability in Xist expression, which may obscure consistent differences at the tissue level. However, these findings are consistent with previous observations that X-chromosome inactivation patterns in PWS patients are largely preserved but exhibit considerable inter-individual variability [62].

Our study has several limitations. First, the conclusions are largely based on correlative observations, as microglial morphology, transcriptional changes, and increased microglia-neuron contacts were not directly linked to functional outcomes such as neuronal activity or behavior. Second, the use of a global *Magel2* knockout model does not fully allow us to distinguish between microglia-intrinsic effects and secondary alterations driven by other cell types or systemic changes. Third, snRNA-seq was performed at a single early developmental time point (P4), limiting insight into temporal dynamics of microglial alterations throughout the lifespan.

In conclusion, our findings identify microglial dysfunction as a potential contributor to hypothalamic circuit abnormalities in PWS, highlighting microglia as a candidate therapeutic target. While pharmacological modulation of microglial activation represents a promising avenue, future studies must establish whether normalizing microglial states can restore neural circuit function and improve metabolic or behavioral outcomes. Given the early-life onset of microglial alterations observed here, timely intervention may be critical. Beyond PWS, these results contribute to a broader understanding of how neuroimmune mechanisms shape neurodevelopmental disorders, with implications for developing targeted therapies that address both neurological and metabolic aspects of disease.

## Animals

Germany: We utilized paternal-allele deleted *Magel2* KO mice [63] at different ages (P4, P10 and 4-5 months), in which the maternally inherited wild-type allele is silent and the paternally derived allele is disrupted. Littermates that inherited the intact maternal allele served as controls. This model has been previously characterized and described in detail in earlier studies [63]. Animals were bred and maintained under standard conditions with *ad libitum* access to food and water, and all experiments were conducted in accordance with institutional animal-care and use guidelines.

United States: All animal experiments were conducted in accordance with the Public Health Service guidelines for the humane care and use of laboratory animals and were approved by the Institutional Animal Care and Use Committee at the University of Michigan (protocol no. PRO00010383). Mice were housed in ventilated cages under controlled temperature conditions (∼23 °C) and a 12-h light/dark cycle (lights on from 6:00 am to 6:00 pm). Animals had ad libitum access to tap water and standard laboratory chow (5L0D; LabDiet), while breeding mice were provided with a breeder chow diet (5008; LabDiet). *Magel2* KO mice were generated by crossing *Magel2* KO male mice with wild-type female mice obtained from The Jackson Laboratory (strain no. 000664). Tail samples collected from postnatal day 1 pups were submitted to TransnetYX for genotyping.

Australia: For Xist RNA FISH experiments, all mice were bred and maintained using standard husbandry procedures under WEHI AEC number 2023.033 and experimental procedures were performed under AEC number 2023.045 with approval from the WEHI animal ethics committee. *Magel2* KO mice were obtained from Jackson Laboratories (Jax stock #009062) and bred on a C57BL/6 background. Animals were housed in an IVC caging system (Techniplast) and maintained in a 14/10 light/dark cycle (lights on 6am-8pm), fed breeder chow.

## Immunohistochemistry

### Mouse

Mice were euthanized using CO and transcardially perfused at a rate of 20 mL/min through the left ventricle with 200 mL of ice-cold 0.01 M PBS, followed by 200 mL of 4% paraformaldehyde (PFA) in 0.3 M PBS (4 °C), while the right atrium was incised to allow drainage. Brains were post-fixed for 24 hours in 4% PFA at 4 °C and subsequently transferred to a 30% sucrose solution (in 0.01 M PBS) at 4 °C for 1–2 days. For immunohistochemical analysis, 50 µm coronal sections were obtained using a Leica Cryostat (CM3050 S) and stored in 0.01 M PBS at 4 °C until processing. Sections were collected in series using the merging anterior commissure as a reference landmark to ensure consistent sampling across animals. Free-floating sections were blocked in 5% normal horse serum in 0.01 M PBS for 1 hour at room temperature, followed by a 15-minute wash in 0.01 M PBS. Sections were then incubated for 24 hours at room temperature in 0.01 M PBS containing 0.1% Triton-X, 0.04% NaN, and the following primary antibodies: IBA1 (1:1000, Fujifilm Wako, rabbit, polyclonal, 019-19741); GFAP (Abcam, goat, ab53554); TUBB3 (1:100, Abcam, rabbit, ab18207); CD68 (Bio-Rad, rat, MCA1957GA); NeuN (Sigma-Aldrich, mouse, MAB377 or Synaptic Systems, guinea pig, 266004); IBA1 (1:200, Synaptic Systems, guinea pig, 234308). After a 15-minute wash in 0.01 M PBS, sections were incubated for 4 hours at room temperature in 0.01 M PBS containing 0.1% Triton-X, 0.04% NaN, and 1:500 Alexa Fluor 488/594/647-conjugated donkey secondary antibodies against rabbit, goat, or mouse (Jackson ImmunoResearch, 711-585-152, 705-585-147, 715-545-151). Finally, sections were washed again for 15 minutes in 0.01 M PBS and mounted with antifade mounting medium containing DAPI (Vectashield, H-1200B).

### Quantification of apoptotic events

Apoptotic cell death was assessed using the NeuroTACS™ Apoptosis Detection Kit (R&D Systems) according to the manufacturer’s instructions. Briefly, two coronal brain sections per animal containing the hypothalamus were processed, including proteinase K treatment, labeling of DNA strand breaks with terminal deoxynucleotidyl transferase (TdT), and detection using a chromogenic substrate resulting in a brown precipitate. Sections were imaged using a conventional light microscope, and apoptotic events were manually quantified based on the presence of brown-stained nuclei. All analyses were performed in a blinded manner.

### PSD95 staining for synaptic engulfment analysis

Three-week-old mice were transcardially perfused with PBS followed by 4% paraformaldehyde (PFA). Brains were dissected and post-fixed in 4% PFA overnight at 4 °C, then cryoprotected by sequential incubation in 10%, 20%, and 30% sucrose solutions for 1 h, 1 h, and overnight, respectively, at 4 °C. Tissues were embedded in optimal cutting temperature compound, stored at −80 °C, and sectioned at 40 µm using a cryostat. For immunostaining, free-floating sections were incubated in blocking solution (3% BSA, 2% goat serum, and 0.1% Triton X-100 in PBS) for 30 min at room temperature, followed by overnight incubation with primary antibodies at 4 °C. After PBS washes, sections were treated with fluorophore-conjugated secondary antibodies (Invitrogen) for 1 h at room temperature. Sections were mounted on SuperFrost slides (Fisher Scientific) and imaged using a Zeiss LSM 780 inverted confocal microscope with ×40 or ×63 objectives. Primary antibodies included PSD95 (1:1,000; Synaptic Systems, catalogue no. 124011) and IBA1 (1:500; Abcam, catalogue no. ab178846).

### Blood-brain barrier perfusion and BBB integrity assessment

Blood–brain barrier (BBB) integrity was assessed by perfusion with fluorescent tracers FITC– dextran (10 kDa), Sigma Aldrich 60842-46-8) and Wheat germ agglutinin (WGA) (40 kDa), ThermoFisher W32466. For labeling, both dyes (0.5 mg/mL each) were added to ice-cold 0.01 M PBS used during transcardial perfusion. Following CO euthanasia, the right atrium was incised and the left ventricle was cannulated for perfusion at a rate of 20 mL/min with 200 mL of dye-containing PBS (4 °C), while protecting samples from light. Immediately after tracer perfusion, fixation was performed with 200 mL of 4% PFA. Analysis of vessel-bound and extravasated dyes was performed via Thresholding and Co-localization via Fiji as described before [26, 64–65].

### Confocal microscopy

Confocal images were acquired using a Nikon Ti2 microscope with AX confocal optics, a Plan Fluor 40x il DIC H N2 objective and a Galvano Bidirectional Scanner. We imaged z-stacks with a step-size of 2 µm and the following settings: Line Averaging: 2x; Resolution: 1024x1024; Dwell Time: 0.5 µsec. Microglial morphology parameters from individual Imaris reconstructions were aggregated per animal and cell type using R 4.4.2. The code used for the Imaris data processing is publicly available at https://github.com/tim-schubert/impro.

### Imaris-assisted three-dimensional reconstruction

Imaris-based analysis of microglial morphology was performed as previously described [26–27, 64]. Slight changes to previous pipelines included: Z-stack images (50 μm depth, 1 μm steps, ×40 magnification) were acquired using a Nikon Ti2 with AX confocal optics. Two image stacks per hemisphere (six sections total) were collected from the center of each sampled area to minimize overlap with neighboring tissue. Raw *.czi* files were processed in Imaris (Version 10.2, Oxford Instruments). Microglial surfaces were reconstructed using custom settings (surface detail 0.500 μm, background subtraction [local contrast], largest fitting sphere 0.7 μm, diffusion transparency 65%). Non-specific background was removed (filter: volume max–400 μm³). Surface reconstructions were then used as templates for process tracing (largest starting-point diameter 7.00 μm, seed points 0.300 μm; removal sphere 15 μm). Seed points were manually corrected when misplaced. Surface and process data were exported to Excel for analysis. For each animal, 8-12 z-stacks were reconstructed. Microglia-neuron interactions were quantified using the surface-to-surface interaction feature, as well as the surface-to-volume ratio function. Appositional and focal interaction points were defined as 1) continuous process-neuron contacts without and 2) locally restricted contacts with substantial thickening of the processes (diameter increase of contact points at the IBA1/TUBB3 intersection >35%), respectively.

### Tissue preparation and library construction for 10x snRNA-sequencing

Postnatal day 4 (P4) pups were euthanized by decapitation, and brains were immediately removed from the skull. The hypothalamus was dissected using a brain matrix, flash-frozen in liquid nitrogen, and stored at −80 °C until processing. Nuclei were isolated as previously described [66]. Briefly, on the day of the experiment, frozen hypothalamic tissue from three mice of the same genotype and sex was pooled and homogenized in lysis buffer (EZ Prep Nuclei Kit, Sigma) supplemented with Protector RNase Inhibitor (Sigma) and 1 M DTT. The homogenate was filtered through a 30 µm MACS strainer (Miltenyi) and centrifuged at 500 rcf for 5 min at 4 °C. Pelleted nuclei were resuspended in wash buffer (10 mM Tris-HCl, pH 8.0; 5 mM KCl; 12.5 mM MgCl; 1% BSA; RNase inhibitor), filtered again, and centrifuged at 500 rcf for 5 min at 4 °C. The washed nuclei were then resuspended in wash buffer containing 7-AAD (BD Biosciences) and subjected to FACS sorting using a MoFlo Astrios Cell Sorter. Sorted nuclei were centrifuged at 500 rcf for 5 min and resuspended in wash buffer. The final nuclei concentration was 7.51 × 10 nuclei/mL. Nuclei samples were submitted to the Advanced Genomics Core at the University of Michigan, where approximately 10,000 nuclei were used to generate snRNAseq libraries using the Chromium Single Cell 3′ Library and Gel Bead Kit v3. Libraries were sequenced on an Illumina NovaSeq 6000 platform (paired-end, 150-nt reads) to a depth of at least 25,000 reads per nucleus.

### Bioinformatic Analysis

Single-nucleus RNA-sequencing (snRNA-seq) data were processed in R(v4.3.3) using the Seurat package(v5.4) [67]. Male and female datasets were analyzed independently throughout the workflow. Raw counts matrices were imported into Seurat objects and subjected to standard preprocessing, including normalization and identification of highly variable features. Data were scaled and integrated across samples using batch-correction strategy to minimize technical variability. Dimensionality reduction was performed using principal component analysis (PCA), and cells were embedded in a low-dimensional space using Uniform Manifold Approximation and Projection (UMAP). Graph-based clustering was performed using Seurat’s shared nearest neighbor approach, and clusters were annotated based on canonical marker gene expression and manual curation. Differential gene expression analysis between *Magel2-KO* and *WT* conditions was performed using a pseudobulk-based approach. Gene expression counts were aggregated per condition using Seurat’s AggregateExpression function [67]. To account for differences in library size, aggregated counts were converted to fractional expression values by normalizing gene counts to the total counts within each condition. Log2 fold changes were then calculated based on the ratio of fractional expression between Magel2-KO and WT samples. Statistical significance was assessed using Fisher’s exact test on aggregated pseudobulk count data for each gene to evaluate proportional differences between conditions. Raw p-values were corrected for multiple testing using the Benjamini-Hochberg false discovery rate procedure. Genes were considered differentially expressed if they met both criteria: adjusted p-value <0.05 and absolute log2 fold change ≥0.25. These results were used as the basis for downstream functional enrichment analyses and visualization. Functional enrichment analyses were performed using the cluster Profiler package(v4.10.1) [68]. Over-representation analysis (ORA) was conducted separately for upregulated and downregulated genes using Gene Ontology (GO) biological process annotations, with enrichment significance assessed using a hypergeometric test and Benjamini-Hochberg correction. Gene set enrichment analysis (GSEA) was performed on ranked gene lists based on log2 fold change using GO biological process, cellular component, and molecular function annotations. Normalized enrichment scores (NES) and adjusted p-values were calculated for each term, enabling comparison of pathway enrichment direction and magnitude between *Magel2* KO and WT conditions.

Integrated pathway heatmaps were generated to summarize cell type-resolved pathway alterations between *Magel2* KO and WT conditions. For each sex, GSEA results from excitatory neurons, inhibitory neurons, astrocytes, OPCs, and microglia were combined into a single matrix. Representative pathways were selected using a direction-balanced strategy, retaining top KO-enriched and WT-enriched GSEA terms for each cell type based on adjusted significance and NES. To enable cross-cell-type comparison, each selected pathway was plotted across all analyzed cell types, rather than only in the cell type from which it was originally selected. Positive NES values indicate pathway enrichment in *Magel2* KO relative to WT, whereas negative NES values indicate enrichment in WT relative to *Magel2* KO. NES values were capped at ±3 for visualization to reduce the influence of extreme values while preserving enrichment directionality. Significantly enriched terms were marked with asterisks using an adjusted q-value threshold of 0.05. Pathways significantly altered in more than one cell type were highlighted to indicate shared or recurrent transcriptional programs. Heatmaps were generated in R using the ComplexHeatmap package(v2.18.0) with a viridis color scale, and matching row and column cell type blocks were outlined to indicate the cell type from which each representative pathway group was selected.

To further characterize microglial functional states, gene modules representing distinct microglial programs (including surveillance, neuroprotection, phagocytosis, inflammation, cytokine production, interferon response, antigen presentation, and proliferation) were defined based on newly published microglia taxonomy [37]. Gene sets were curated directly from this study and applied to the dataset to quantify program activity at the single-cell level. Module scoring was performed using rank-based approaches implemented in R, enabling comparison of functional state distributions between Magel2-KO and WT conditions across sexes.

Cell-cell communication analysis was performed using the CellChat package (v1.6.1) [44]. For each sex, the integrated Seurat object was split into WT and Magel2-KO conditions and analyzed independently. CellChat inference was conducted using mouse ligand-receptor interaction database. Overexpressed genes and ligand-receptor pairs were identified, and communication probabilities were computed using the triMean method with correction for cell population size. Interactions supported by fewer than 10 cells were excluded. Communication networks were aggregated at both interaction and pathway levels, and pathway-specific information flow was calculated to quantify signaling activity. Comparative analyses between conditions included assessment of interaction number, interaction strength, and pathway-level signaling differences, as well as cell-type-specific incoming and outgoing communication patterns. All analyses were performed in R, and figures were generated using ggplot2(v4.0.2) (Wickham, 2016) and package-specific visualization functions.

### Statistical analysis and visualization

All statistical analyses were performed in R. Multiple testing correction was applied using the Benjamini–Hochberg method where appropriate. Data visualization included UMAP embeddings, volcano plots, heatmaps, barplots, dot plots, and network diagrams. Figures were generated using ggplot2 and related visualization packages, with publication-quality formatting.

### Xist analysis

Hypothalamus was collected from *Magel2* KO pups and wildtype littermates at postnatal day 4, washed in PBS and incubated in 1x Trypsin/EDTA for 15 minutes at 37°C. Tissue was then washed in PBS before being mechanically dissociated, first with a wide-bore pipette tip followed by trituration with a standard tip. The cell suspension was passed through a 70 µm MACS® SmartStrainer (Miltenyi Biotec) into a 15ml polystyrene round-bottomed tube and the filter was flushed into the same tube with PBS. Cell suspensions were centrifuged and the pellet was resuspended in PBS containing 0.5% BSA before being cytospun onto coverslips. Xist RNA-FISH performed on cytospun cells as described [69] with minor adaptations. In brief, Xist RNA was detected with the 15kb cDNA, pCMV-Xist-PA [70]. 1μg Xist cDNA was used in a nick translation reaction (Abbott) to generate DNA probes labelled with SEEBRIGHT® Orange 552 dUTP (Enzo Life Sciences). ∼100ng probe per sample was precipitated in Ethanol with 10% NaOAc, and 1μg Salmon Sperm (Life Technologies) followed by resuspension in formamide (Sigma-Aldrich). Probes were then denatured at 75°C for 10 minutes and kept on ice for a maximum of 30 minutes during cell preparation. Coverslips containing cytospun cells were fixed in 4% (w/v) paraformaldehyde made in nuclease-free PBS for 10 minutes at room temperature and then permeabilised on ice for 5 minutes in cold PBS containing 0.5% (v/v) Triton X-100 and 2mM Ribonucleoside Vanadyl Complex (RVC). Coverslips were washed thrice in 70% (v/v) Ethanol for 5 minutes and then dehydrated through 5 minute washes in 80% (v/v), 95% (v/v) and 100% Ethanol sequentially. An equal volume of 2X hybridisation buffer [4X SSC (Sigma-Aldrich), 40% (w/v) dextran sulfate (Sigma-Aldrich), 2mg/ml BSA (New England Biolabs), and 400mM RVC] was added to denatured probe in formamide and aliquoted onto a microscope slide. Coverslips were then placed cell-side-down on the probe droplet and incubated overnight at 37°C in a humidified chamber (humidified with 50% formamide, 50% 2X SSC) for hybridisation. The following morning coverslips were washed thrice for 5 minutes in 50% formamide, 50% 2X SSC at 42°C and then a further 3 times in 2X SSC at 42°C. Coverslips were counterstained with 0.2µg/ml DAPI nuclear stain (ThermoFisher) in 2X SSC at room temperature for 1 minute followed by 2 quick washes in 2X SSC before mounting on microscope slides in VECTASHIELD® Vibrance™ antifade mounting medium (Vector Laboratories). Images were captured using an LSM 880 Upright (Zeiss) confocal microscope, analysis was done using the open-source ImageJ distribution package FIJI and manual scoring.

## Human studies

### Subject information

Thirteen post-mortem hypothalami were obtained from the Netherlands Brain Bank (NBB), and consent was obtained from donors or their legal guardians for brain autopsy and use of medical records and brain tissue for research purposes. Five PWS cases and eight matched controls were included. PWS diagnosis was confirmed by Multiplex Ligation-dependent Probe Amplification-based genotyping, demonstrating copy number alterations consistent with PWS. Specifically, PWS subjects showed a 50% gene dosage reduction in the BP2– BP3 region of chromosome 15q11-q13 region (including MKRN3 and MAGEL2). Controls were defined as individuals without PWS-associated genomic deletions and without known endocrine, metabolic, or neurological disorders. Groups were matched for sex, age, tissue fixation time (FT), and post-mortem delay (PMD).

| Group | Age | Females<br>(n; %) | PMD<br>(minutes) | FT<br>(days) |
| --- | --- | --- | --- | --- |
| Control (n=8) | 44.63 ± 13.41 | 4; 50 | 854.20 ± 750.10 | 64.25 ± 33.70 |
| PWS (n=5) | 34.00 ± 9.46 | 3; 60 | 1388.00 ± 1312.00 | 38.67 ± 17.79 |
| p-value | 0.152 | 1 | 0.405 | 0.206 |

Values are demonstrated as mean ± standard deviation. T test was employed for age, PMD and FT, and Exact Fisher test for biological sex.

### Immunofluorescence

After brain autopsy, hypothalamic tissue was immersed in 10% phosphate-buffered formalin at room temperature. Brain tissue was ethanol-dehydrated, toluene-cleared, and paraffin-embedded. All the formalin-fixed paraffin-embedded (FFPE) hypothalamic tissue was coronally serially sectioned from rostral to caudal at 6 μm. Anatomical orientation of the paraventricular nucleus of the hypothalamus was determined by Nissl staining, also referencing well-defined anatomical landmarks (e.g., fornix), and posteriorly confirmed by the presence of arginine-vasopressin-containing neurons (data not shown). The section with the highest number of neuronal soma was used for microglial staining. FFPE hypothalamic sections were deparaffinized in 100% xylene, rehydrated in grading ethanol (100% −50%), and rinsed in distilled water. Sections were washed in 0.05 M Tris buffered saline (TBS). Heat induced epitope retrieval using microwave treatment (10 min 700 W) was performed with Tris buffer (pH 9.0). After cooling, sections were incubated with 3% hydrogen peroxide in SUMI incubation buffer (0.25% gelatin, 0.5% Triton X 100 in 0.05 M TBS) for 10 minutes to eliminate endogenous peroxidase activity. Sections were then washed in TBS and incubated with primary antibodies for 1 h at room temperature and then overnight at 4°C. Next, sections were rinsed and incubated with biotinylated secondary antibodies. C1q signal was enhanced using a biotin-conjugated antibody. For immunofluorescence, the appropriate fluorescent secondary antibody was employed and DAPI counterstaining was performed. Images were acquired using a Leica TCS SP8 confocal microscope (Leica Microsystems, Wetzlar, Germany) in 32-bit sequential standard mode at 400 Hz, with a resolution of 1024 × 1024 pixels, objective lens 40X 3–5 fields/regions of interest per sample labeled with IBA1 and C1q. Variation between fields reflects inherent tissue characteristics, and hypothalamic paraventricular nucleus topography. The laser intensity was constant within the image and defined in a pilot acquisition.

### Statistical analysis

All statistical analyses were performed in R (v4.4.2) or GraphPad Prism (v10.6.0). Data are presented as mean ± SEM unless otherwise indicated. Normality was assessed using the Shapiro-Wilk test together with visual inspection of QQ plots and residual distributions. Equality of variances was evaluated using F tests or Levene’s test where appropriate. For comparisons between two groups, two-tailed unpaired Welch’s t-tests were used unless stated otherwise. For experiments involving multiple factors (e.g., genotype × distance in Sholl analysis), two-way ANOVA was performed followed by Tukey’s post hoc multiple-comparisons test. Categorical variables were analyzed using chi-square tests. Correlation analyses were performed using linear regression and Pearson correlation coefficients. For imaging-based analyses, biological replicates corresponded to individual animals or human subjects, whereas technical replicates included multiple sections, z-stacks, or fields of view acquired per subject. Quantifications from individual cells or image stacks were averaged per animal before statistical testing unless otherwise specified. Investigators were blinded to genotype and condition during image acquisition, reconstruction, and quantification. No statistical methods were used to predetermine sample size, but sample sizes were comparable to previous studies employing similar methodologies. No samples or animals were excluded unless tissue quality or staining artifacts prevented reliable analysis. For snRNA-seq analyses, male and female datasets were analyzed independently. Differential expression analysis was performed using pseudobulk aggregation of counts per condition and cell type. P values were adjusted for multiple testing using the Benjamini-Hochberg false discovery rate (FDR) correction. Genes with adjusted p < 0.05 and absolute log2 fold change ≥ 0.25 were considered differentially expressed. Gene Ontology over-representation analysis and GSEA were corrected using Benjamini-Hochberg adjustment. CellChat analyses excluded inferred interactions supported by fewer than 10 cells, and communication probabilities were computed using the triMean method with population-size correction. Exact statistical tests, sample sizes, and adjusted p values are reported in the corresponding figure legends. Statistical significance was defined as p < 0.05.

## AI-assisted language editing

During manuscript preparation, ChatGPT (OpenAI) was used to assist with proofreading, grammar correction, language refinement, and improvement of readability. All scientific content, data interpretation, analyses, and conclusions were developed and verified by the authors.

## Data availability

Raw data will be available from the corresponding author upon reasonable request.

## Acknowledgements

The authors thank Jeffrey M. Friedman for resources and assistance with experiments involving *Magel2* KO mice. We also thank the Nikon Imaging Center at BioQuant, Heidelberg, for technical support and access to imaging facilities.

## Supplementary Figures

**Supplementary Figure 1.**
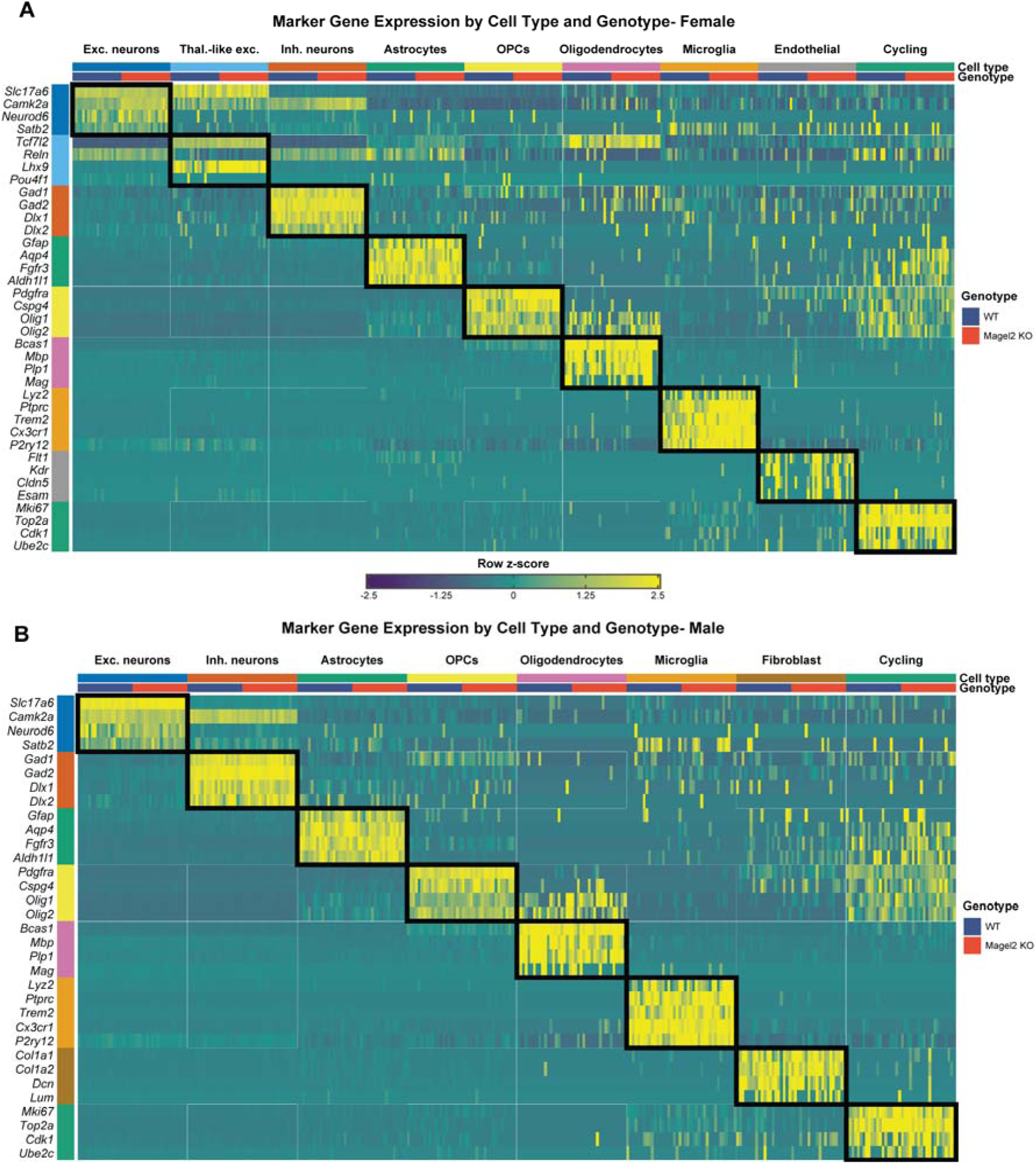
Canonical marker gene expression supports cell-type annotation across sex and genotype. (A-B) Binned marker gene expression heatmaps across annotated hypothalamic cell types and genotypes in female (A) and male (B) P4 snRNA-seq datasets. Rows represent canonical marker genes grouped by their expected cell-type identity, and columns represent binned nuclei grouped by cell type and genotype. For each cell type-genotype group, nuclei were randomly partitioned into 20 bins, and average normalized expression was calculated per bin to reduce single-nucleus-level sparsity while preserving cell-type and genotype structure. Expression values are shown as row-scaled z-scores capped at ±2.5. Top annotations indicate cell type and genotype, with WT and *Magel2* KO shown separately within each cell-type block. Left-side row annotations indicate marker-gene groups. Black boxes highlight the expected correspondence between each marker-gene group and its matching annotated cell type. Canonical neuronal, glial, vascular, mesenchymal, immune, and cycling-cell markers show selective enrichment in their corresponding annotated populations, supporting the robustness of cell-type annotation across both female and male datasets and demonstrating that major cell identities are preserved across genotypes.

**Supplementary Figure 2.**
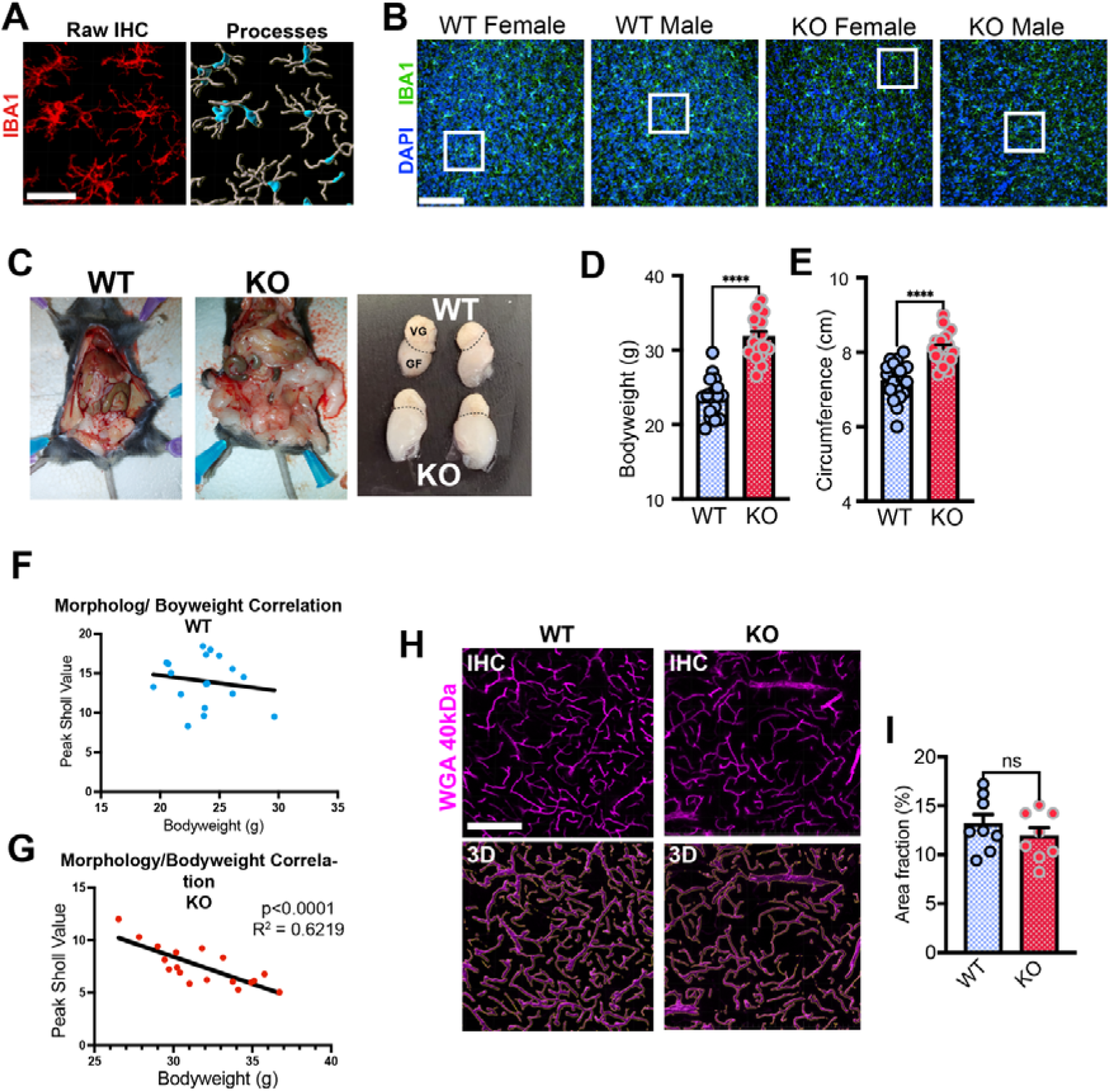
**A** Example three-dimensional reconstruction of microglial processes via Imaris. Confocal images with raw IBA1 signal are used as a template for subsequent surface and process reconstruction. Scale bar = 25 µm. **B** Overview confocal images showing DAPI and NeuN staining in the hypothalamus of female and male WT and *Magel2* KO mice. White boxes correspond with the high magnification examples depicted in Figure 1. Scale bar = 100 µm. **C** Typical examples of fat tissue and gonadal fat pads 4-month-old male WT and Magel2 KO mice. Note the excessive visceral fat in the *Magel2* KO mouse. Dashed lines separate gonadal fat pads (GF) from vesicular glands (VG). **D** *Magel2* KO mice display markedly increased bodyweight at 5 months of age, n=18 per group (9 male / 9 female). **E** Abdominal circumference is significantly increased in 6-month-old *Magel2* KO mice. **F** Correlation of microglial morphology (peak Sholl value) with bodyweight in WT mice, n=18 (9 male / 9 female). **G** Correlative analysis shows that a loss of microglial complexity is associated with increased body weight in *Magel2* KO mice. **H, I** Assessment of vascularization via transcardiac perfusion with WGA 40 kDa to label cerebral blood vessels, n=8 per group (4 male / 4 female) Scale bar = 100 µm.

**Supplementary Figure 3.**
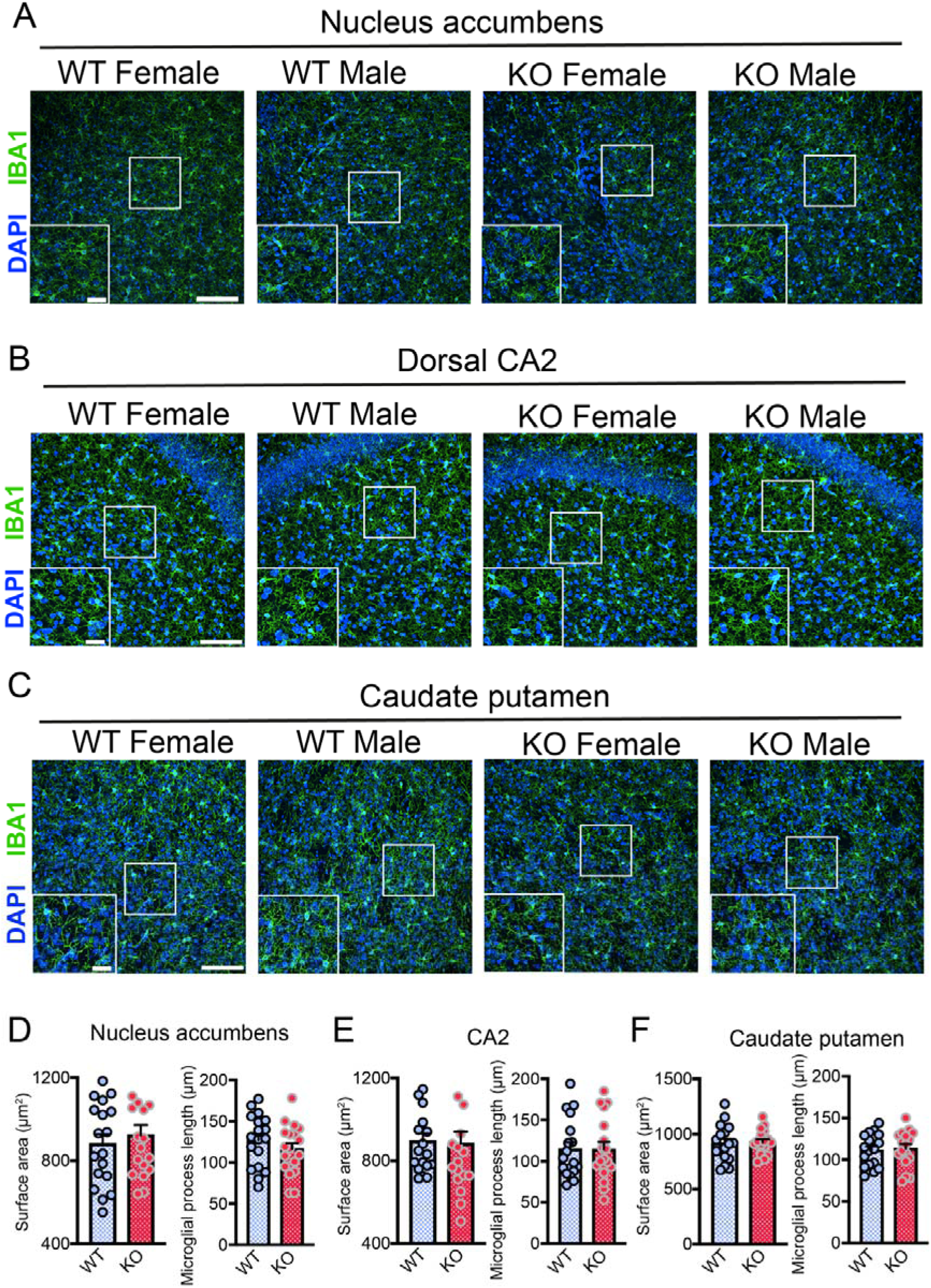
**A** Overview confocal images highlight microglial distribution in the NAc of WT and *Magel2 KO* mice. Enlarged insets highlight detailed microglial morphology in different experimental subgroups. Scale bars = 10 µm and 50 µm. **B** Overview confocal images highlight microglial distribution in the CA2 of WT and *Magel2 KO* mice. Enlarged insets highlight detailed microglial morphology in different experimental subgroups. Scale bars = 10 µm and 50 µm. **C** Overview confocal images highlight microglial distribution in the caudate putamen of WT and *Magel2 KO* mice. Enlarged insets highlight detailed microglial morphology in different experimental subgroups. Scale bars = 10 µm and 50 µm. **D** Quantification of surface area and process length in the NAc of WT and *Magel2* KO mice, n=18 per group (9 males, 9 females). **E** Quantification of surface area and process length in the CA2 of WT and *Magel2* KO mice, n=18 per group (9 males, 9 females). **F** Quantification of surface area and process length in the caudate putamen of WT and *Magel2* KO mice, n=18 per group (9 males, 9 females).

**Supplementary Figure 4.**
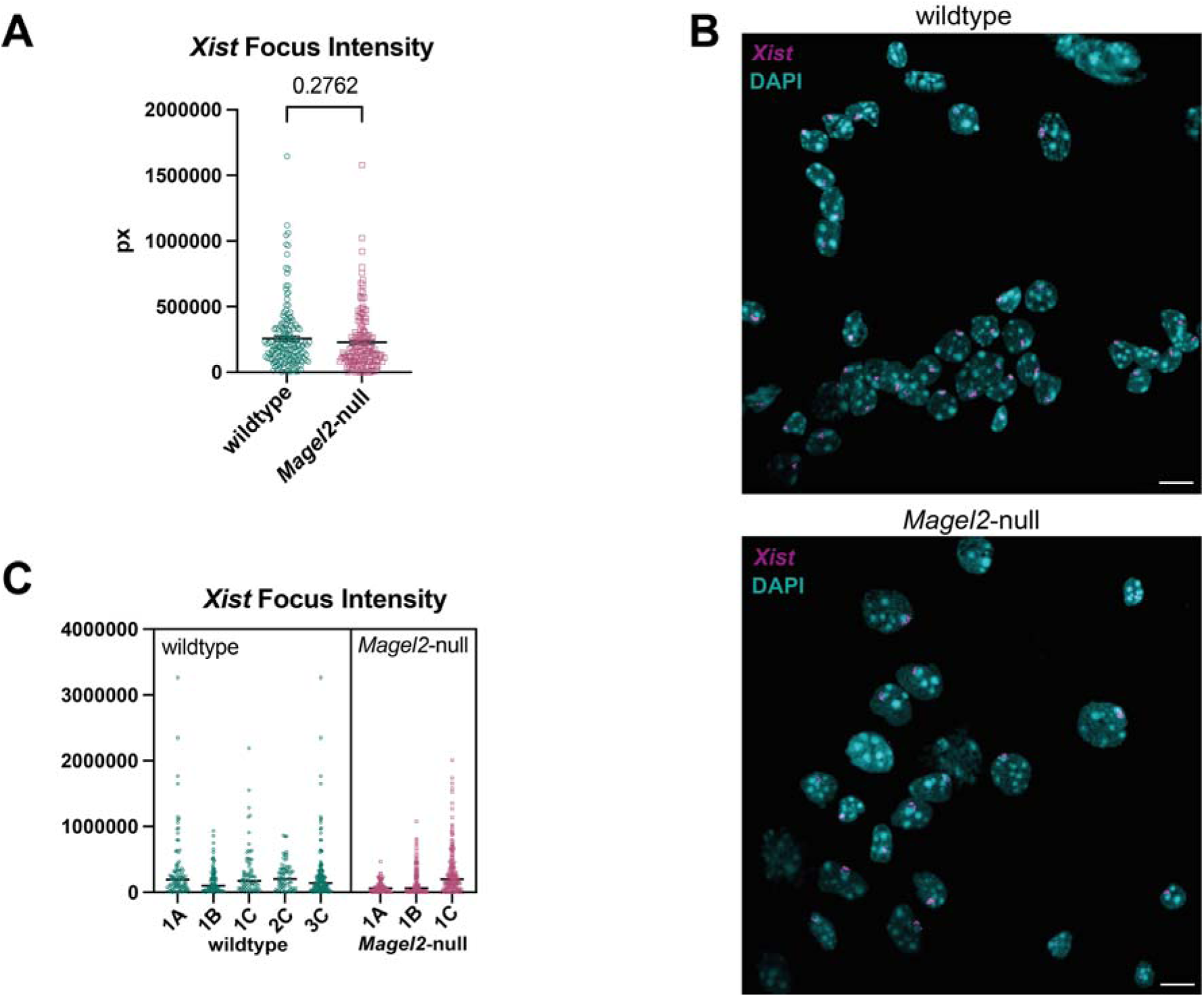
**A** Graph plotting the total fluorescence intensity for each *Xist* focus, n >340 nuclei scored per genotype from wildtype (n=5) and *Magel2* KO (n=3) mice. **B** Representative images from wildtype (top) and *Magel2* KO (bottom) showing RNA-FISH staining for *Xist* (magenta), nuclei stained with DAPI (cyan), scale bar 10 µm. **C** Nested data displaying biological variance between individuals, letters represent litter matching n >70 nuclei scored per mouse.

**Supplementary Figure 5.**
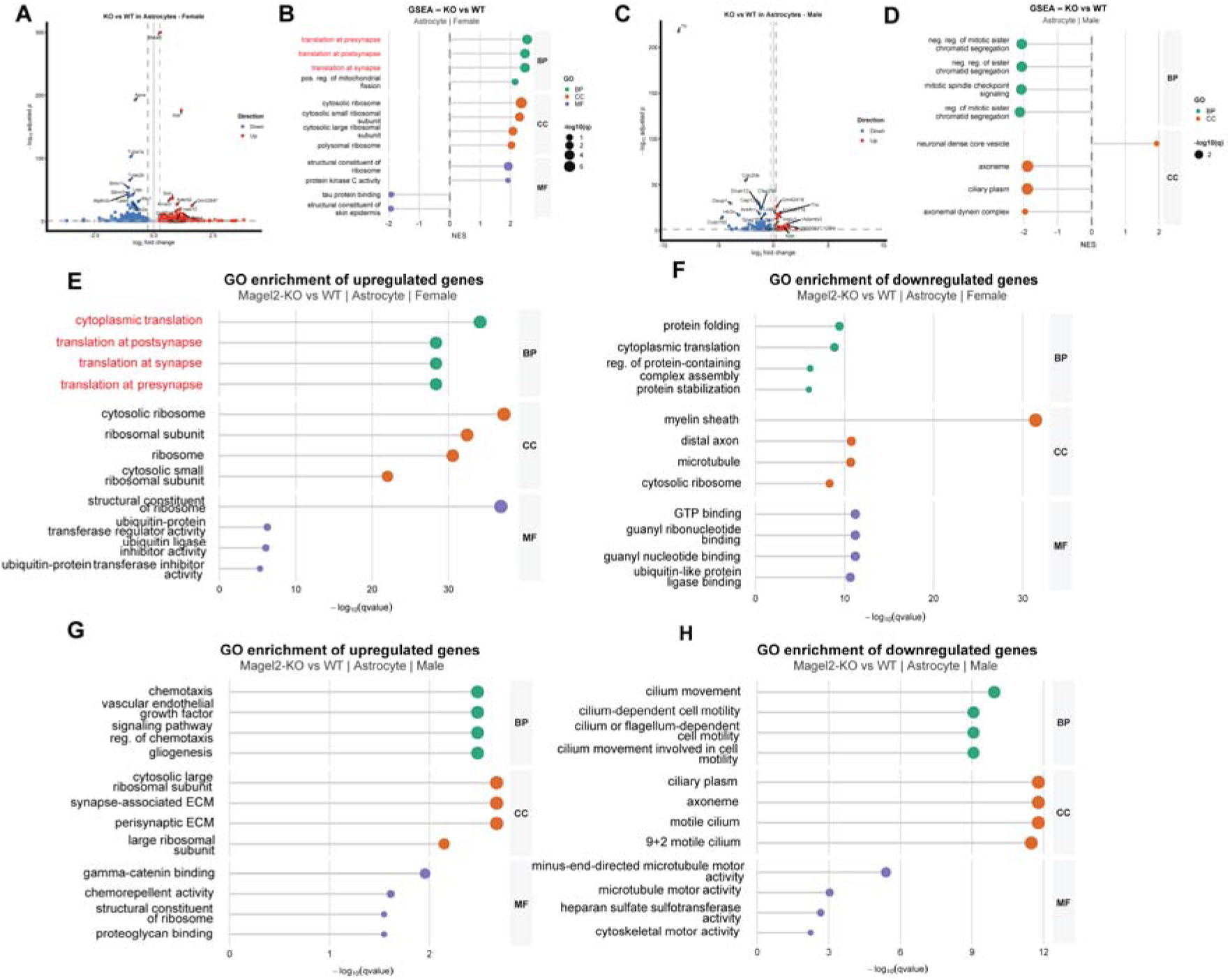
Magel2-dependent transcriptional and pathway alterations in astrocytes. (A) Volcano plot showing differential gene expression in astrocytes from female mice comparing *Magel2* KO and wild-type (WT) conditions. Significantly upregulated and downregulated genes are defined by |log fold change| > 0.5 and p < 0.05. (B) Gene set enrichment analysis (GSEA) of astrocytes from female samples based on ranked log fold changes, highlighting enriched biological pathways in *Magel2* KO relative to WT. (C) Volcano plot of differential gene expression in astrocytes from male mice (Magel2-KO vs WT). (D) GSEA results for male astrocytes. (E–F) Gene Ontology (GO) over-representation analysis (ORA) of differentially expressed genes in female astrocytes. (E) Downregulated pathways in *Magel2* KO; (F) upregulated pathways. (G–H) GO ORA of astrocyte DEGs in male samples showing (G) downregulated and (H) upregulated pathways in *Magel2* KO mice.

**Supplementary Figure 6.**
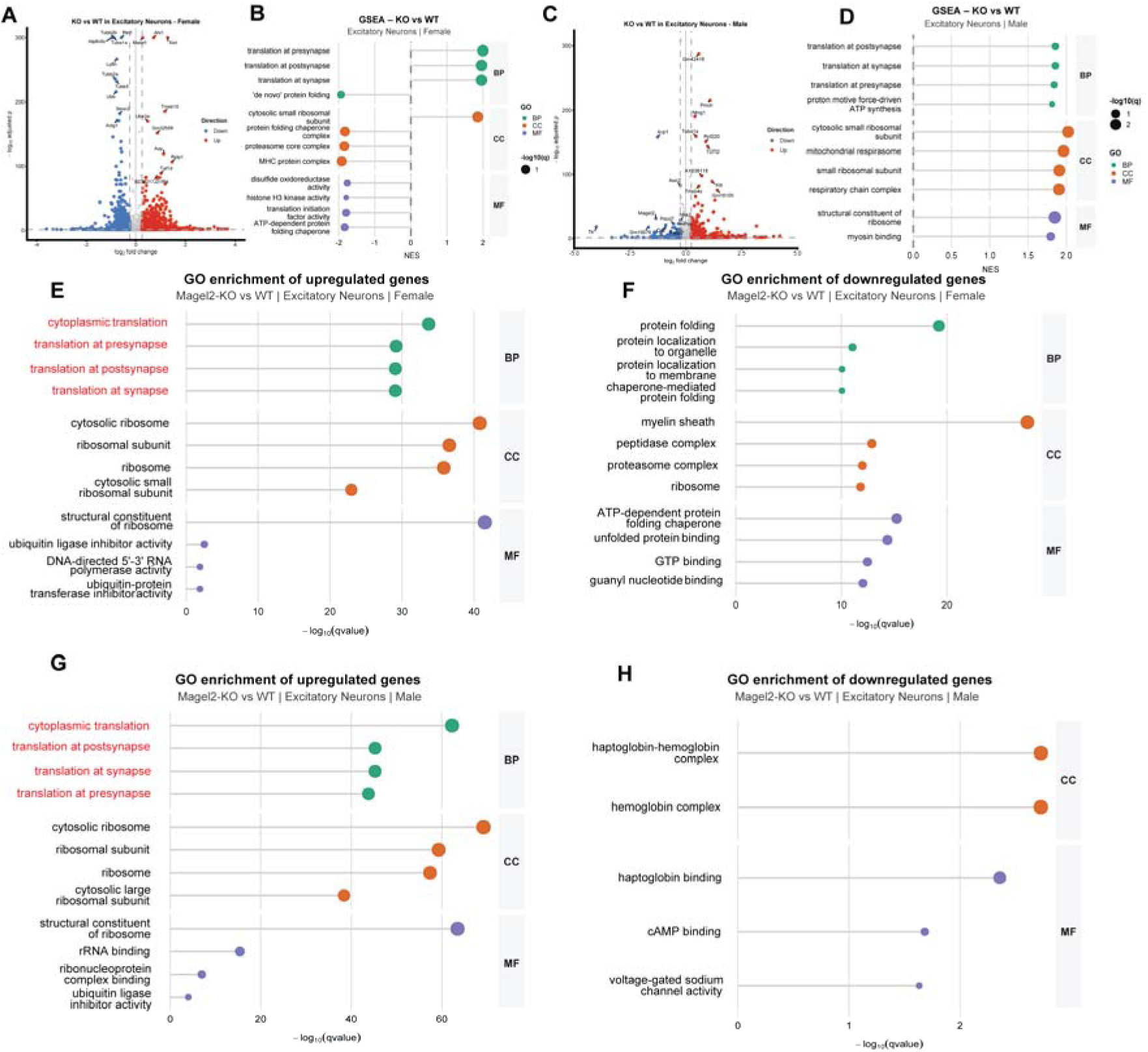
*Magel2* deficiency alters transcriptional programs in excitatory neurons. (A) Volcano plot showing differential gene expression in excitatory neurons from female mice comparing *Magel2* KO and wild-type (WT) conditions. Significantly upregulated and downregulated genes are defined by |log fold change| > 0.5 and p < 0.05. (B) Gene set enrichment analysis (GSEA) of excitatory neurons from female samples based on ranked log fold changes. (C) Volcano plot of differential gene expression in excitatory neurons from male mice (Magel2-KO vs WT). (D) GSEA results for male excitatory neurons. (E–F) Gene Ontology (GO) over-representation analysis (ORA) of differentially expressed genes in female excitatory neurons. (E) Downregulated pathways in *Magel2* KO; (F) upregulated pathways. (G–H) GO ORA of excitatory neuron DEGs in male samples showing (G) downregulated and (H) upregulated pathways in *Magel2* KO mice.

**Supplementary Figure 7.**
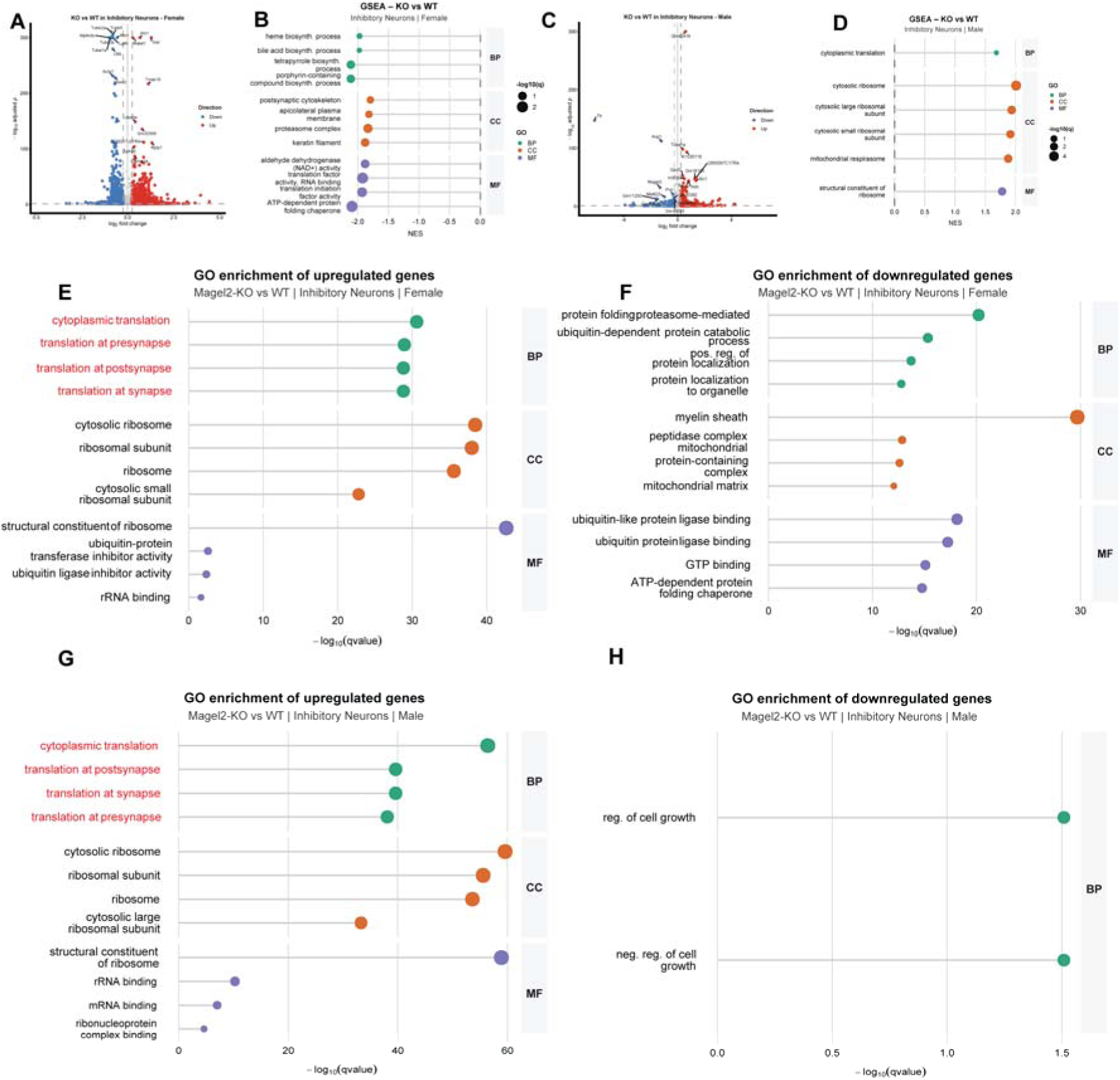
Transcriptional and pathway alterations in inhibitory neurons following *Magel2* loss. (A) Volcano plot showing differential gene expression in inhibitory neurons from female mice comparing *Magel2* KO and wild-type (WT) conditions. Significantly upregulated and downregulated genes are defined by |log fold change| > 0.5 and p < 0.05. (B) Gene set enrichment analysis (GSEA) of inhibitory neurons from female samples based on ranked log fold changes. (C) Volcano plot of differential gene expression in inhibitory neurons from male mice (*Magel2* KO vs WT). (D) GSEA results for male inhibitory neurons. (E–F) Gene Ontology (GO) over-representation analysis (ORA) of differentially expressed genes in female inhibitory neurons. (E) Downregulated pathways in *Magel2* KO; (F) upregulated pathways. (G–H) GO ORA of inhibitory neuron DEGs in male samples showing (G) downregulated and (H) upregulated pathways in *Magel2* KO mice.

**Supplementary Figure 8.**
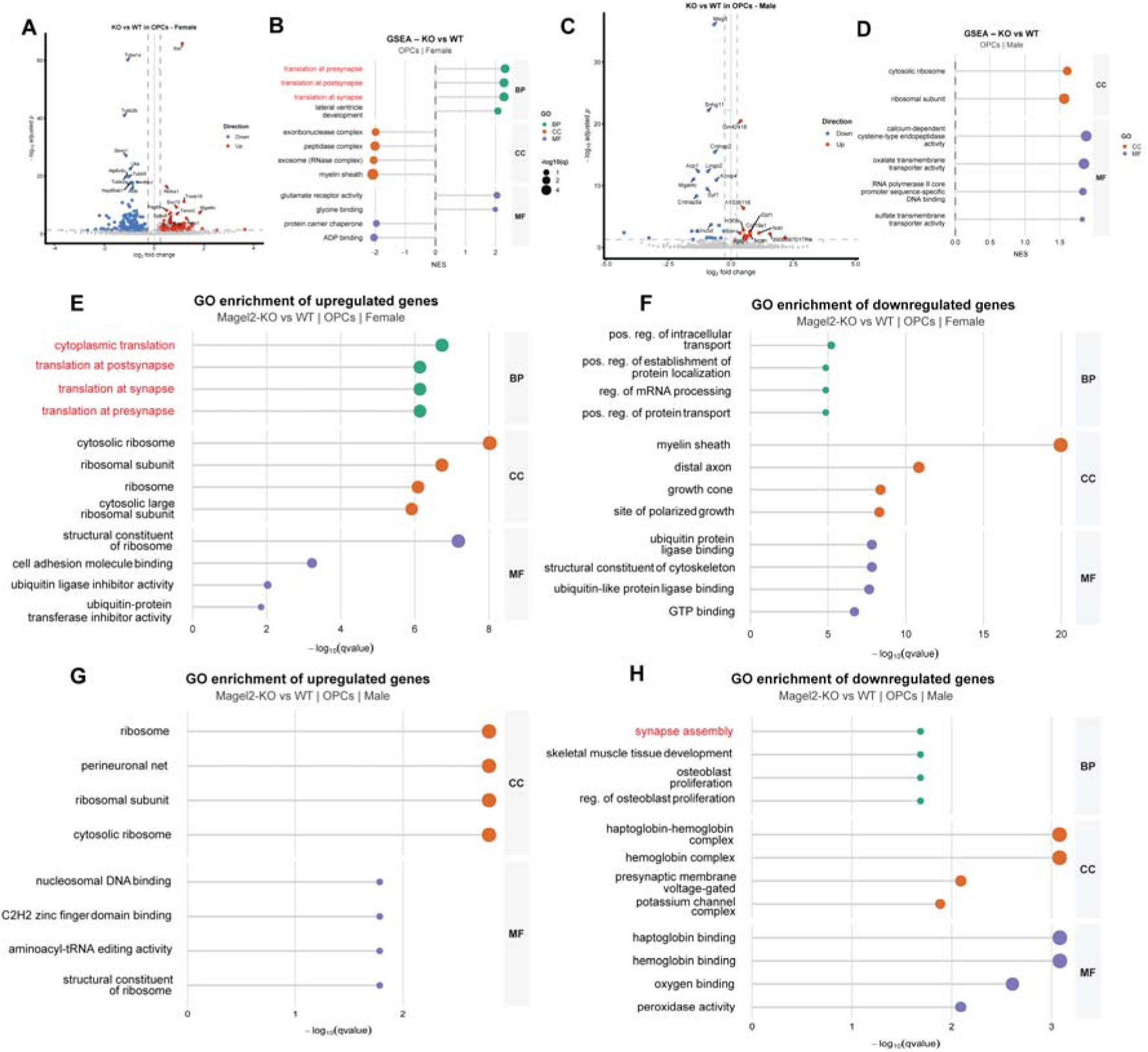
*Magel2* deficiency induces transcriptional and functional changes in oligodendrocyte precursor cells (OPCs)1. (A) Volcano plot showing differential gene expression in OPCs from female mice comparing *Magel2* KO and wild-type (WT) conditions. Significantly upregulated and downregulated genes are defined by |log fold change| > 0.5 and p < 0.05. (B) Gene set enrichment analysis (GSEA) of OPCs from female samples based on ranked log fold changes. (C) Volcano plot of differential gene expression in OPCs from male mice (*Magel2* KO vs WT). (D) GSEA results for male OPCs. (E–F) Gene Ontology (GO) over-representation analysis (ORA) of differentially expressed genes in female OPCs. (E) Downregulated pathways in *Magel2* KO; (F) upregulated pathways. (G–H) GO ORA of OPC DEGs in male samples showing (G) downregulated and (H) upregulated pathways in *Magel2* KO mice.

**Supplementary Figure 9.**
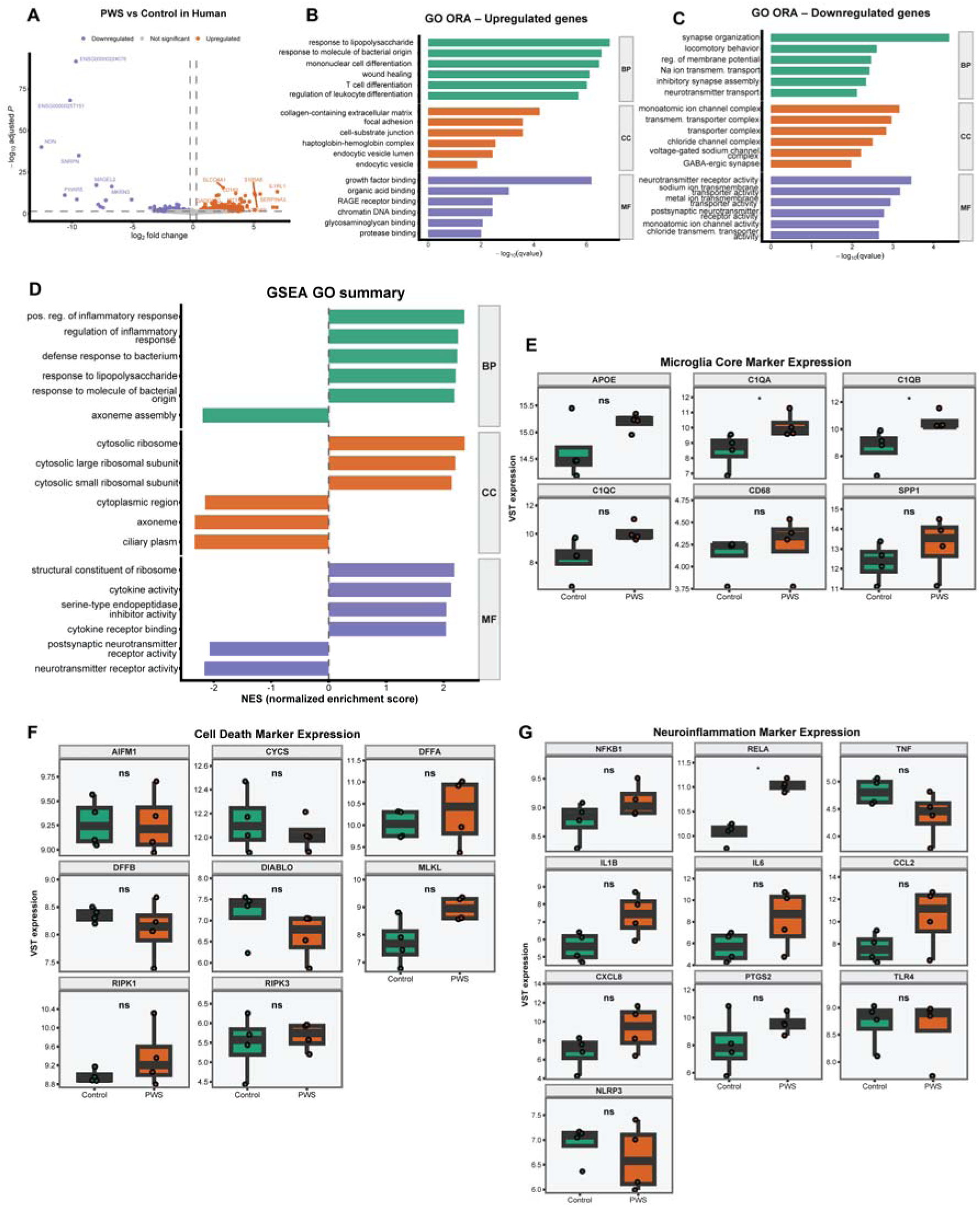
Bulk RNA-seq analysis of human PWS hypothalamic samples reveals immune activation and synaptic dysregulation. **(A)** Volcano plot showing differential gene expression between Prader–Willi syndrome (PWS) and healthy control human samples. Differential expression was performed using DESeq2. Genes with FDR < 0.05 and |log2FC| ≥ 0.2 are highlighted (orange: upregulated in PWS; blue: downregulated in PWS). **(B)** Gene Ontology (GO) over-representation analysis (ORA) of upregulated genes in PWS. Enriched terms are grouped by ontology categories: Biological Process (BP), Cellular Component (CC), and Molecular Function (MF). Bar length represents statistical significance (−log10(q-value)). Enrichment highlights immune-related processes, including response to lipopolysaccharide, leukocyte differentiation, and wound healing, as well as extracellular matrix and vesicle-associated components. **(C)** GO ORA of downregulated genes in PWS. Enriched terms are predominantly associated with synaptic organization, neurotransmitter transport, and ion channel activity, indicating impaired neuronal signaling and synaptic function. **(D)** Gene Set Enrichment Analysis (GSEA) summarizing pathway-level changes between PWS and control samples. Enriched pathways include immune and stress-related processes, alongside depletion of neuronal and synaptic signaling programs. **(E)** Expression levels of core microglial marker genes (*APOE, C1QA, C1QB, C1QC, CD68, SPP1*) across conditions, confirming the presence and transcriptional modulation of microglial signatures in bulk tissue. **(F)** Expression analysis of cell death–related genes (*AIFM1, CYCS, DFFA, DFFB, DIABLO, MLKL, RIPK1, RIPK3*), indicating altered apoptotic and survival pathways in PWS samples. **(G)** Expression of neuroinflammation-associated genes (*NFKB1, RELA, TNF, IL1B, IL6, CCL2, CXCL8, PTGS2, TLR4, NLRP3*), demonstrating increased inflammatory signaling in PWS compared to controls.

