## Supplementary Information for "*Magel2* deficiency drives microglial synaptic engulfment and alters neuron-microglia interactions in Prader-Willi syndrome"

Melbourne, Australia

^5^ Department of Medical Biology, University of Melbourne, Melbourne, Australia

^6^ Institute of Cell and Gene Technology, Shenzhen University of Advanced Technology, Shenzhen, China

^7^ Bioinformatics and Omics Data Analytics, DKFZ, Heidelberg, Germany

^8^ National Center for Tumor Diseases (NCT) Heidelberg

^9^ Department of Animal Sciences, The Ohio State University, Columbus, OH, United States

^10^ Department of Molecular and Integrative Physiology, University of Michigan Medical School, United States

^11^ Institute  of  Pathology,  Heidelberg  University  Hospital, Heidelberg, Germany.

^12^ Translational Lung Research Center (TLRC) Heidelberg, member of the DZL, Heidelberg, Germany

^13^ Blizard Institute, Barts and The London School of Medicine and Dentistry, Queen Mary University of London, London, UK

^14^ Department of Medical Physiology, College of Medicine, Texas A&M University, Bryan, TX, USA

^15^ Institute for Translational Metabolism Research, Helmholtz Diabetes Center at Helmholtz Munich, Neuherberg, Germany

^16^ German Center for Diabetes Research (DZD), 85764 Neuherberg, Germany

^17^ University of Cambridge, Metabolic Research Laboratories and NIHR Cambridge Biomedical Research Centre, Addenbrookes Hospital, Cambridge, United Kingdom

^18^ Institut de Neurobiologie de la Méditerranée (INMED), INSERM, Aix-Marseille Université, 13273 Marseille, France

^19^ Department of Anatomy and Neurobiology, University of California, Irvine, Irvine, CA 92617, USA

* These authors contributed equally

**Supplementary Figures**

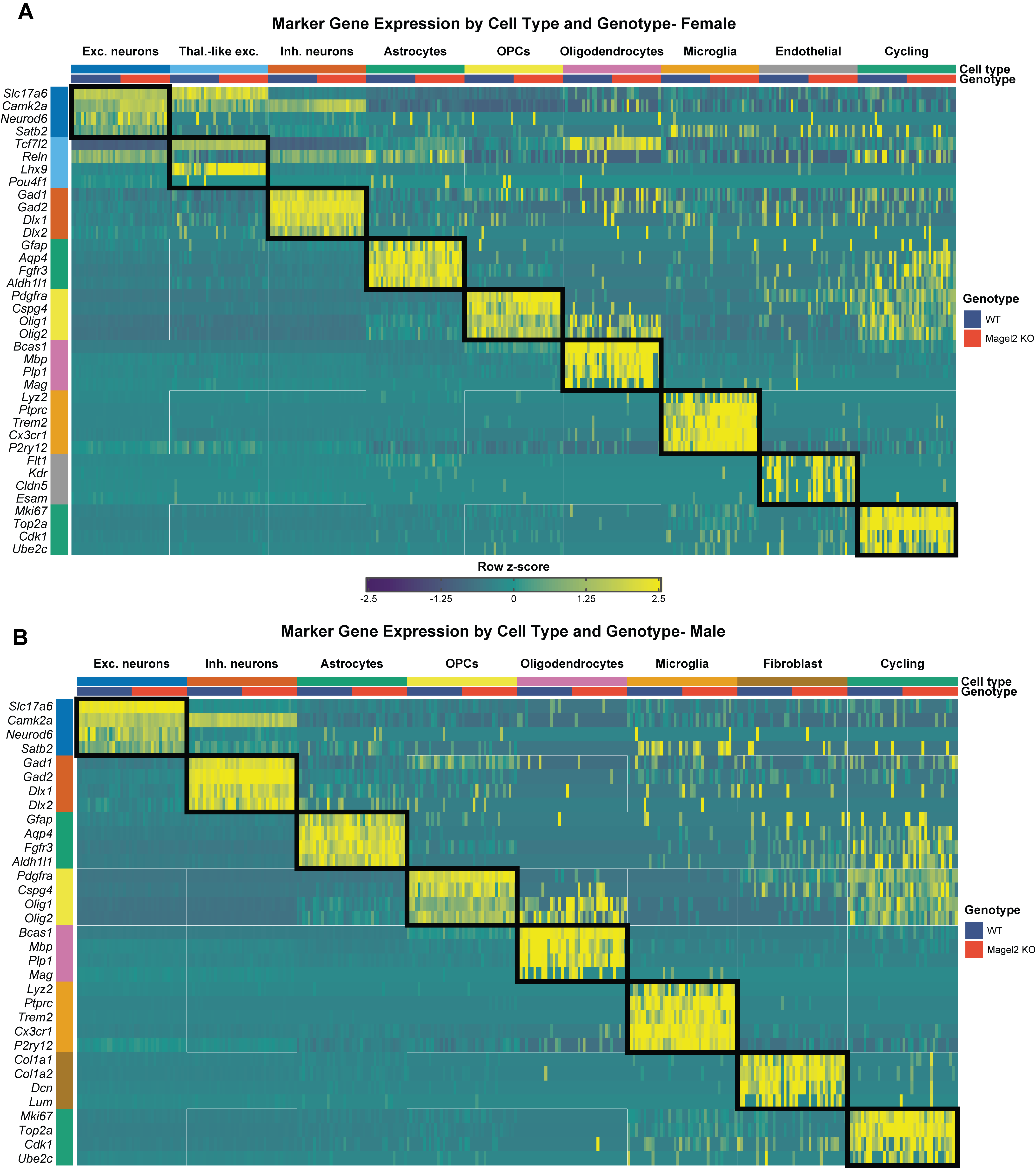

**Supplementary Figure 1**

**Canonical marker gene expression supports cell-type annotation across sex and genotype.** (A-B) Binned marker gene expression heatmaps across annotated hypothalamic cell types and genotypes in female (A) and male (B) P4 snRNA-seq datasets. Rows represent canonical marker genes grouped by their expected cell-type identity, and columns represent binned nuclei grouped by cell type and genotype. For each cell type-genotype group, nuclei were randomly partitioned into 20 bins, and average normalized expression was calculated per bin to reduce single-nucleus-level sparsity while preserving cell-type and genotype structure. Expression values are shown as row-scaled z-scores capped at ±2.5. Top annotations indicate cell type and genotype, with WT and *Magel2* KO shown separately within each cell-type block. Left-side row annotations indicate marker-gene groups. Black boxes highlight the expected correspondence between each marker-gene group and its matching annotated cell type. Canonical neuronal, glial, vascular, mesenchymal, immune, and cycling-cell markers show selective enrichment in their corresponding annotated populations, supporting the robustness of cell-type annotation across both female and male datasets and demonstrating that major cell identities are preserved across genotypes.

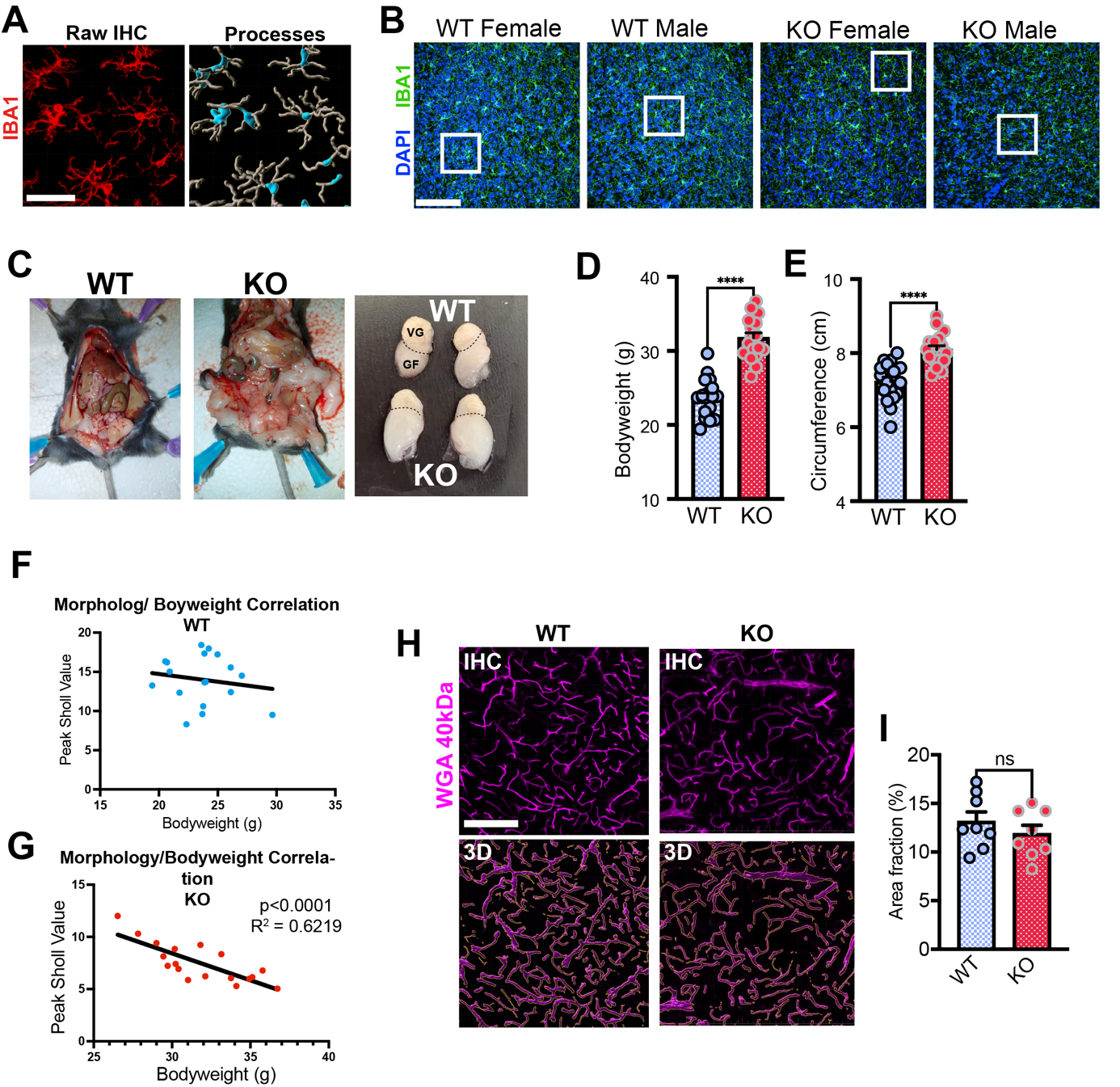

**Supplementary Figure 2**

**A** Example three-dimensional reconstruction of microglial processes via Imaris. Confocal images with raw IBA1 signal are used as a template for subsequent surface and process reconstruction. Scale bar = 25 µm. **B** Overview confocal images showing DAPI and NeuN staining in the hypothalamus of female and male WT and *Magel2* KO mice. White boxes correspond with the high magnification examples depicted in Figure 1. Scale bar = 100 µm. **C** Typical examples of fat tissue and gonadal fat pads 4-month-old male WT and Magel2 KO mice. Note the excessive visceral fat in the *Magel2* KO mouse. Dashed lines separate gonadal fat pads (GF) from vesicular glands (VG). **D** *Magel2* KO mice display markedly increased bodyweight at 5 months of age, n=18 per group (9 male / 9 female). **E** Abdominal circumference is significantly increased in 6-month-old *Magel2* KO mice. **F** Correlation of microglial morphology (peak Sholl value) with bodyweight in WT mice, n=18 (9 male / 9 female). **G** Correlative analysis shows that a loss of microglial complexity is associated with increased body weight in *Magel2* KO mice. **H, I** Assessment of vascularization via transcardiac perfusion with WGA 40 kDa to label cerebral blood vessels, n=8 per group (4 male / 4 female) Scale bar = 100 µm.

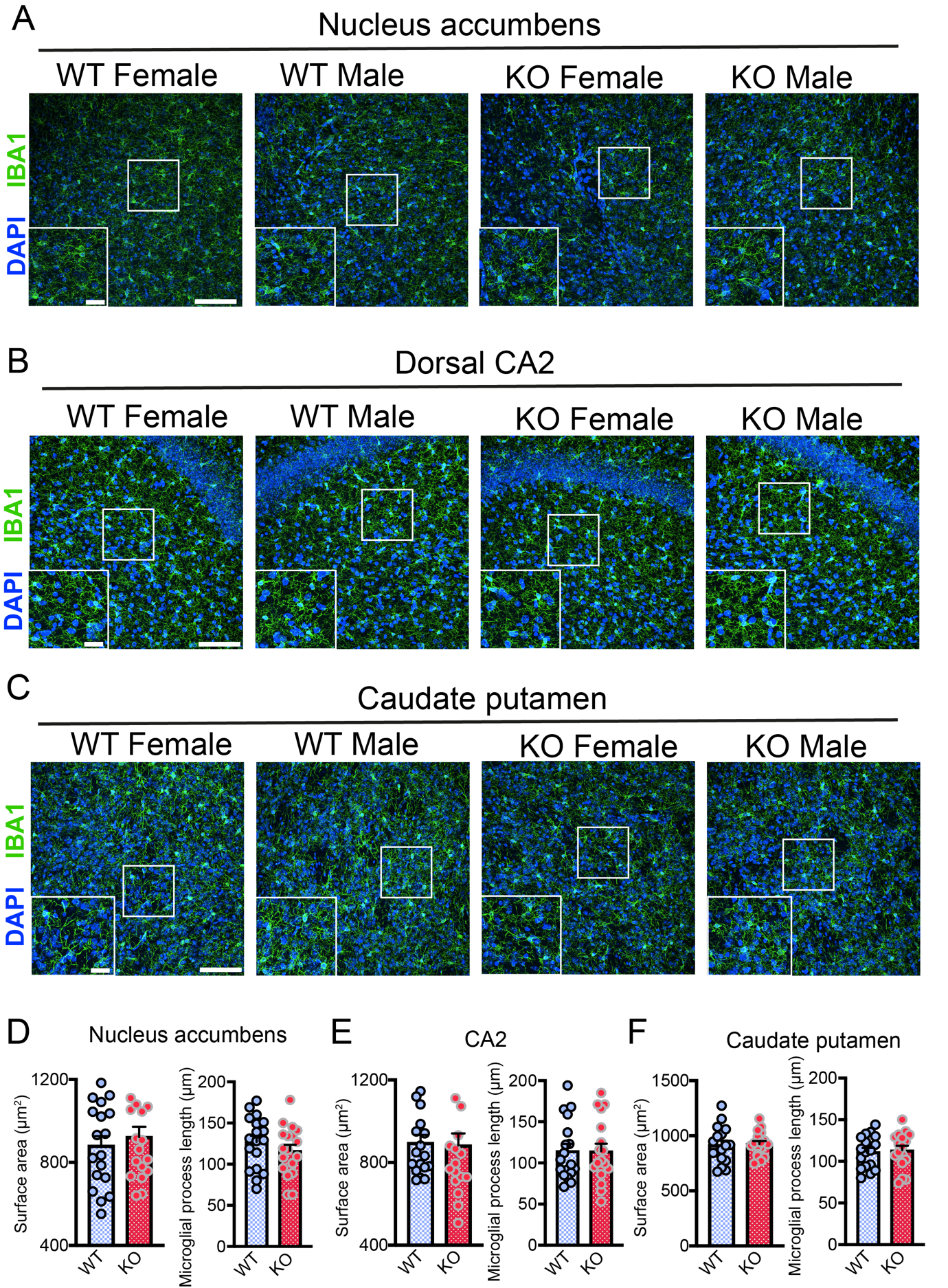

**Supplementary Figure 3**

**A** Overview confocal images highlight microglial distribution in the NAc of WT and *Magel2 KO* mice. Enlarged insets highlight detailed microglial morphology in different experimental subgroups. Scale bars = 10 µm and 50 µm. **B** Overview confocal images highlight microglial distribution in the CA2 of WT and *Magel2 KO* mice. Enlarged insets highlight detailed microglial morphology in different experimental subgroups. Scale bars = 10 µm and 50 µm. **C** Overview confocal images highlight microglial distribution in the caudate putamen of WT and *Magel2 KO* mice. Enlarged insets highlight detailed microglial morphology in different experimental subgroups. Scale bars = 10 µm and 50 µm. **D** Quantification of surface area and process length in the NAc of WT and *Magel2* KO mice, n=18 per group (9 males, 9 females). **E** Quantification of surface area and process length in the CA2 of WT and *Magel2* KO mice, n=18 per group (9 males, 9 females). **F** Quantification of surface area and process length in the caudate putamen of WT and *Magel2* KO mice, n=18 per group (9 males, 9 females).

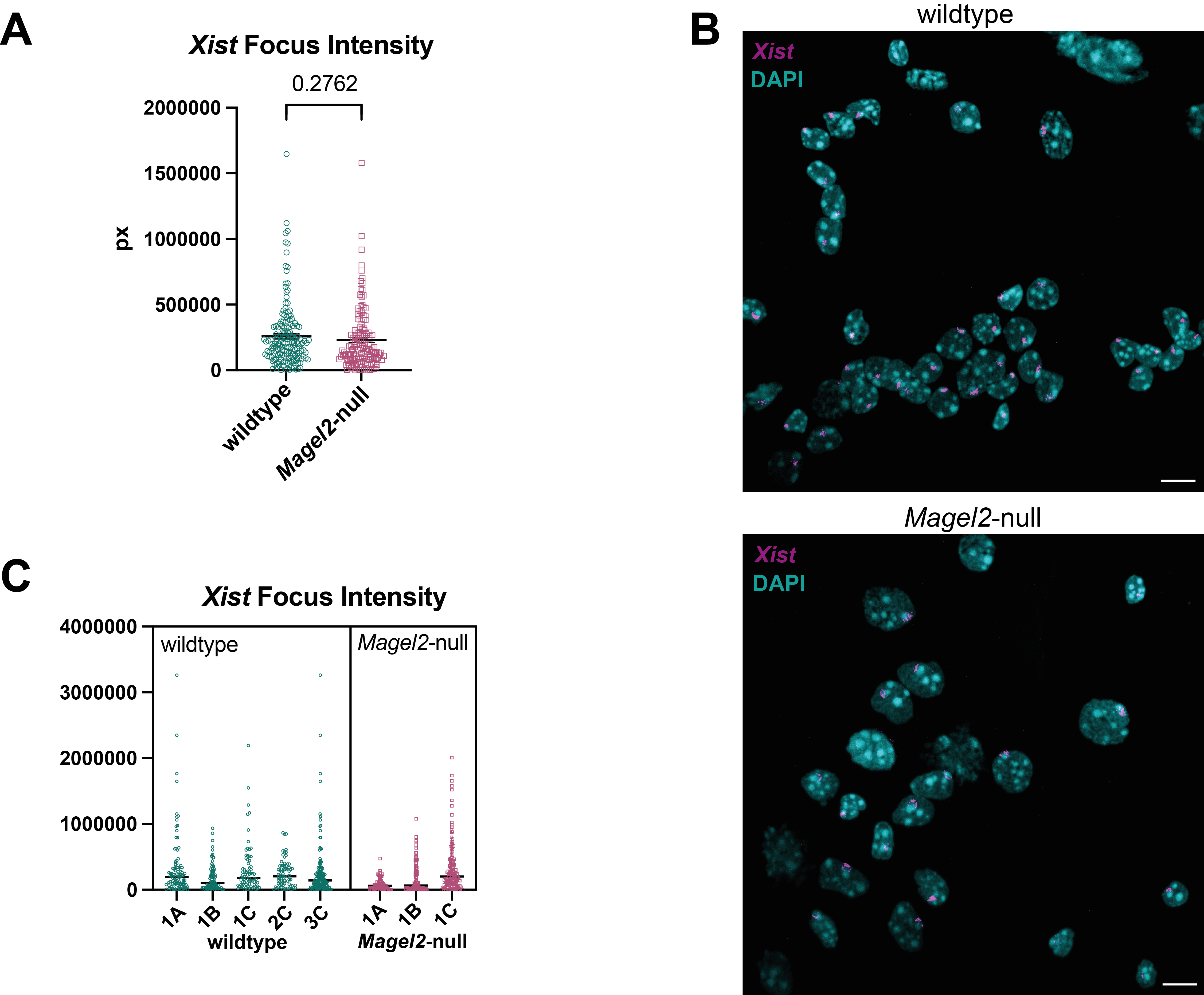

**Supplementary Figure 4**

**A** Graph plotting the total fluorescence intensity for each *Xist* focus, n >340 nuclei scored per genotype from wildtype (n=5) and *Magel2* KO (n=3) mice. **B** Representative images from wildtype (top) and *Magel2* KO (bottom) showing RNA-FISH staining for *Xist* (magenta), nuclei stained with DAPI (cyan), scale bar 10 µm. **C** Nested data displaying biological variance between individuals, letters represent litter matching n >70 nuclei scored per mouse.

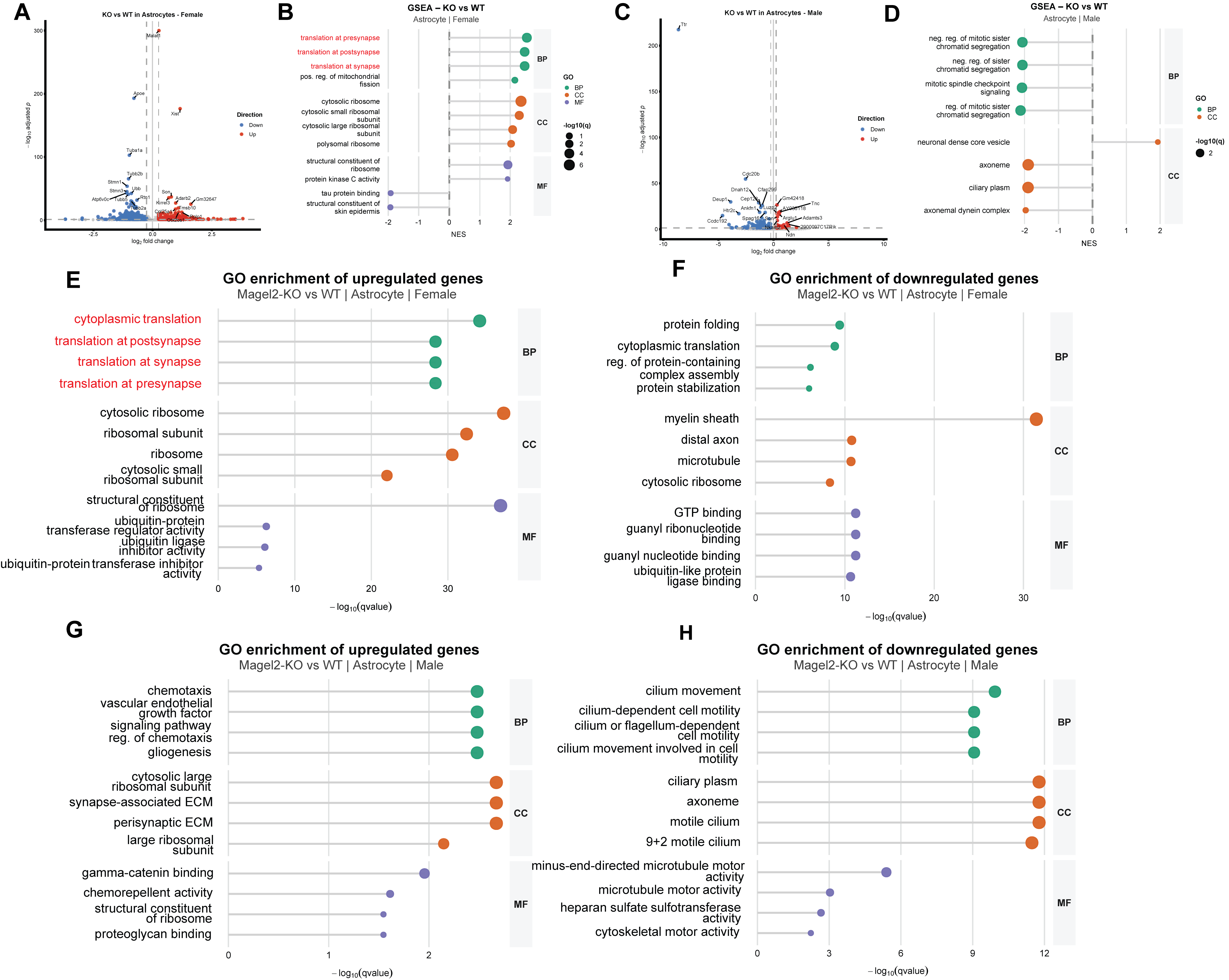

**Supplementary Figure 5**

**Magel2-dependent transcriptional and pathway alterations in astrocytes**
(A) Volcano plot showing differential gene expression in astrocytes from female mice comparing *Magel2* KO and wild-type (WT) conditions. Significantly upregulated and downregulated genes are defined by |log₂ fold change| > 0.5 and p < 0.05.
(B) Gene set enrichment analysis (GSEA) of astrocytes from female samples based on ranked log₂ fold changes, highlighting enriched biological pathways in *Magel2* KO relative to WT.
(C) Volcano plot of differential gene expression in astrocytes from male mice (Magel2-KO vs WT). (D) GSEA results for male astrocytes. (E–F) Gene Ontology (GO) over-representation analysis (ORA) of differentially expressed genes in female astrocytes. (E) Downregulated pathways in *Magel2* KO; (F) upregulated pathways.
(G–H) GO ORA of astrocyte DEGs in male samples showing (G) downregulated and (H) upregulated pathways in *Magel2* KO mice.

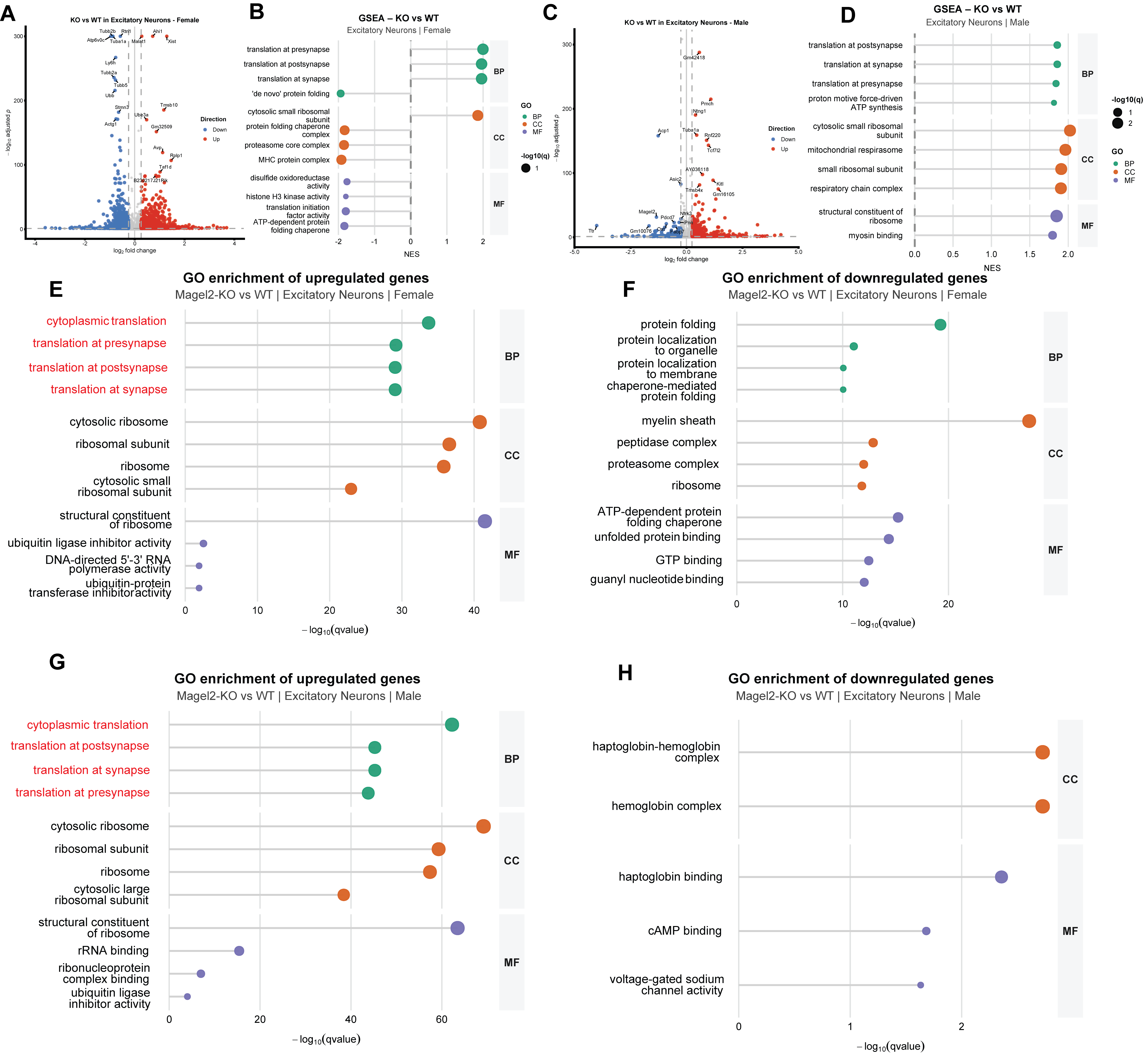

**Supplementary Figure 6**

***Magel2* deficiency alters transcriptional programs in excitatory neurons**
(A) Volcano plot showing differential gene expression in excitatory neurons from female mice comparing *Magel2* KO and wild-type (WT) conditions. Significantly upregulated and downregulated genes are defined by |log₂ fold change| > 0.5 and p < 0.05.
(B) Gene set enrichment analysis (GSEA) of excitatory neurons from female samples based on ranked log₂ fold changes. (C) Volcano plot of differential gene expression in excitatory neurons from male mice (Magel2-KO vs WT). (D) GSEA results for male excitatory neurons.
(E–F) Gene Ontology (GO) over-representation analysis (ORA) of differentially expressed genes in female excitatory neurons. (E) Downregulated pathways in *Magel2* KO; (F) upregulated pathways. (G–H) GO ORA of excitatory neuron DEGs in male samples showing (G) downregulated and (H) upregulated pathways in *Magel2* KO mice.

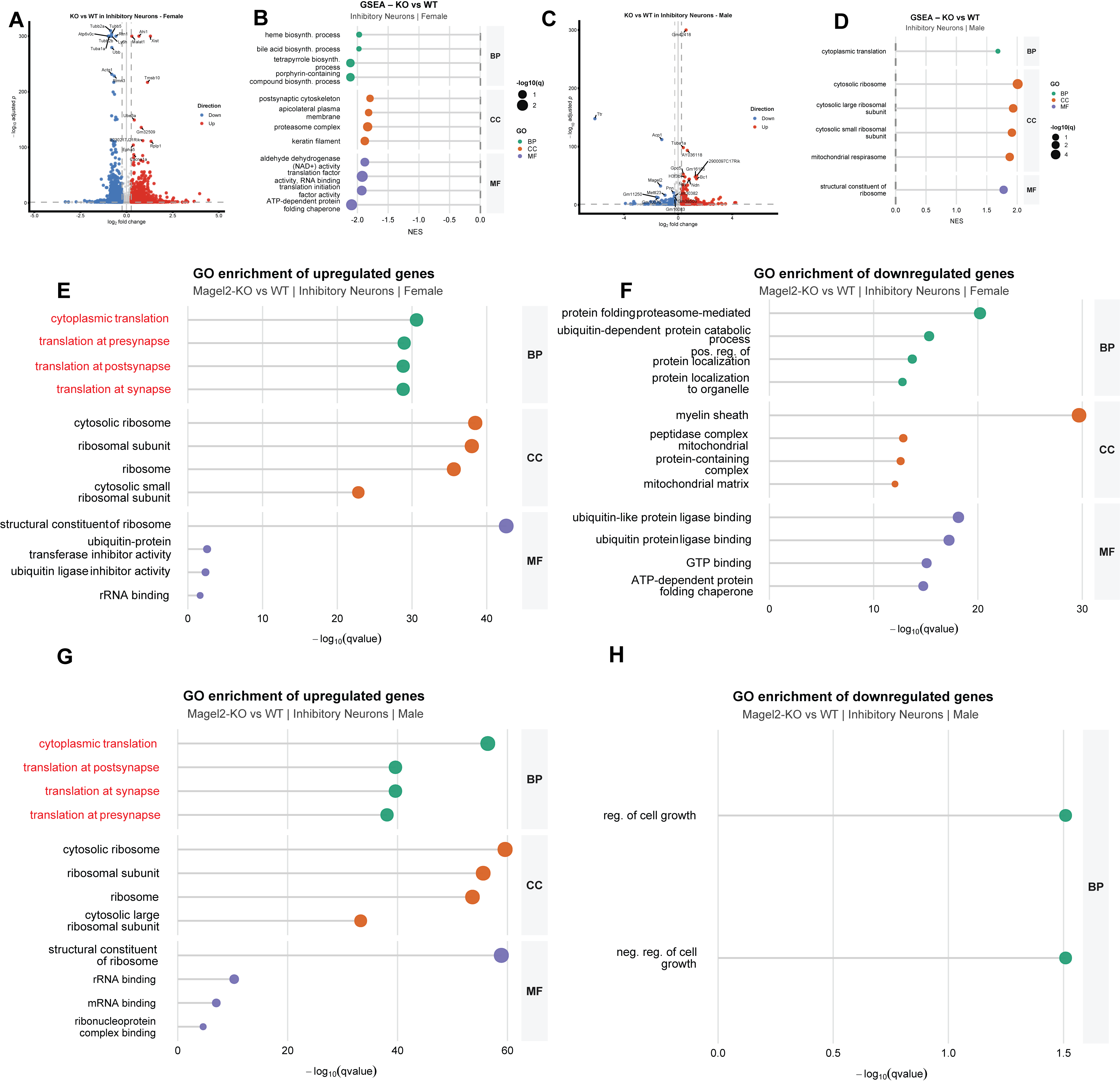

**Supplementary Figure 7**

**Transcriptional and pathway alterations in inhibitory neurons following *Magel2* loss**

(A) Volcano plot showing differential gene expression in inhibitory neurons from female mice comparing *Magel2* KO and wild-type (WT) conditions. Significantly upregulated and downregulated genes are defined by |log₂ fold change| > 0.5 and p < 0.05.
(B) Gene set enrichment analysis (GSEA) of inhibitory neurons from female samples based on ranked log₂ fold changes. (C) Volcano plot of differential gene expression in inhibitory neurons from male mice (*Magel2* KO vs WT). (D) GSEA results for male inhibitory neurons.
(E–F) Gene Ontology (GO) over-representation analysis (ORA) of differentially expressed genes in female inhibitory neurons. (E) Downregulated pathways in *Magel2* KO; (F) upregulated pathways. (G–H) GO ORA of inhibitory neuron DEGs in male samples showing (G) downregulated and (H) upregulated pathways in *Magel2* KO mice.

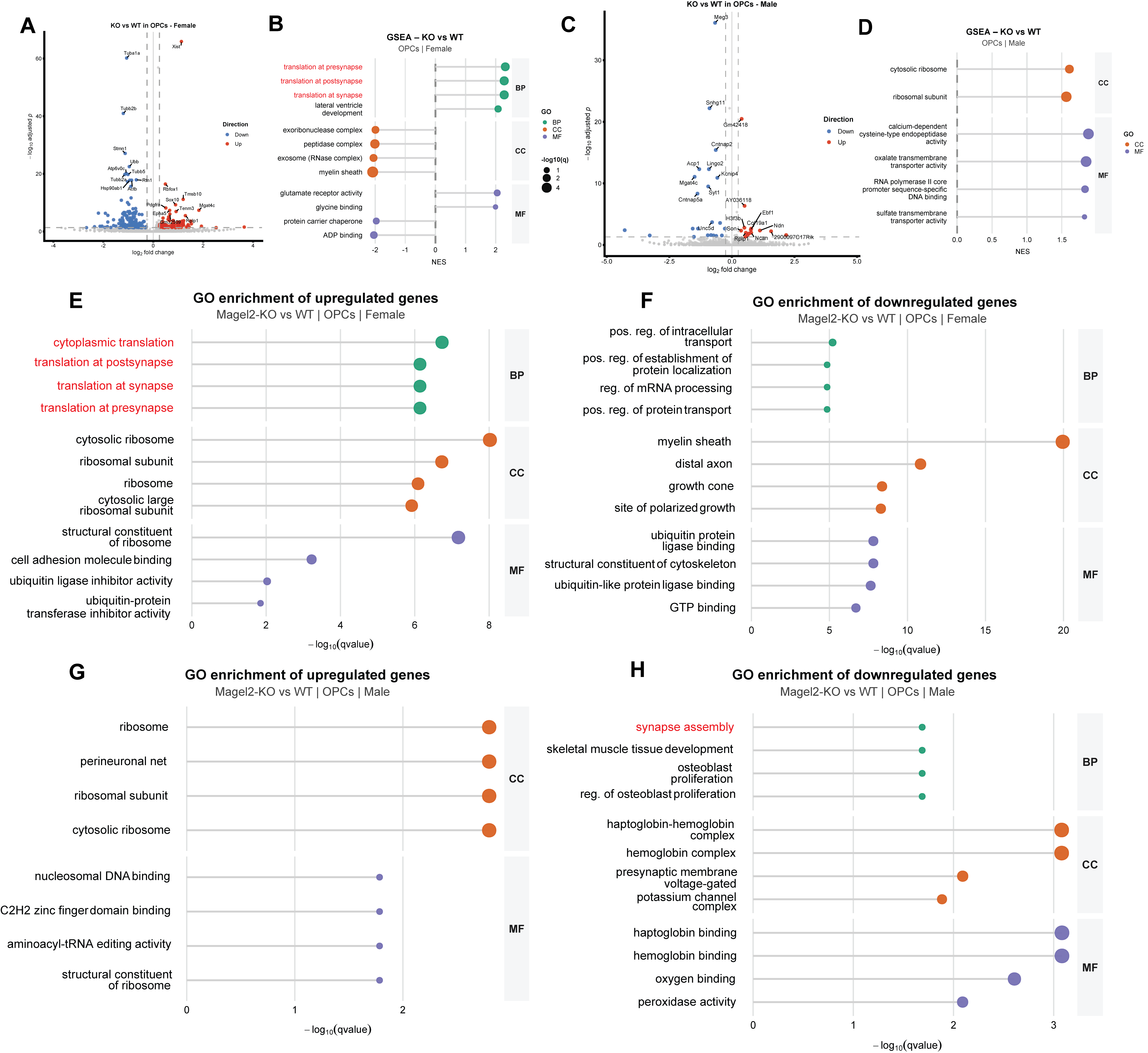

**Supplementary Figure 8**

***Magel2* deficiency induces transcriptional and functional changes in oligodendrocyte precursor cells (OPCs)**

(A) Volcano plot showing differential gene expression in OPCs from female mice comparing *Magel2* KO and wild-type (WT) conditions. Significantly upregulated and downregulated genes are defined by |log₂ fold change| > 0.5 and p < 0.05.
(B) Gene set enrichment analysis (GSEA) of OPCs from female samples based on ranked log₂ fold changes. (C) Volcano plot of differential gene expression in OPCs from male mice (*Magel2* KO vs WT). (D) GSEA results for male OPCs. (E–F) Gene Ontology (GO) over-representation analysis (ORA) of differentially expressed genes in female OPCs. (E) Downregulated pathways in *Magel2* KO; (F) upregulated pathways.
(G–H) GO ORA of OPC DEGs in male samples showing (G) downregulated and (H) upregulated pathways in *Magel2* KO mice.

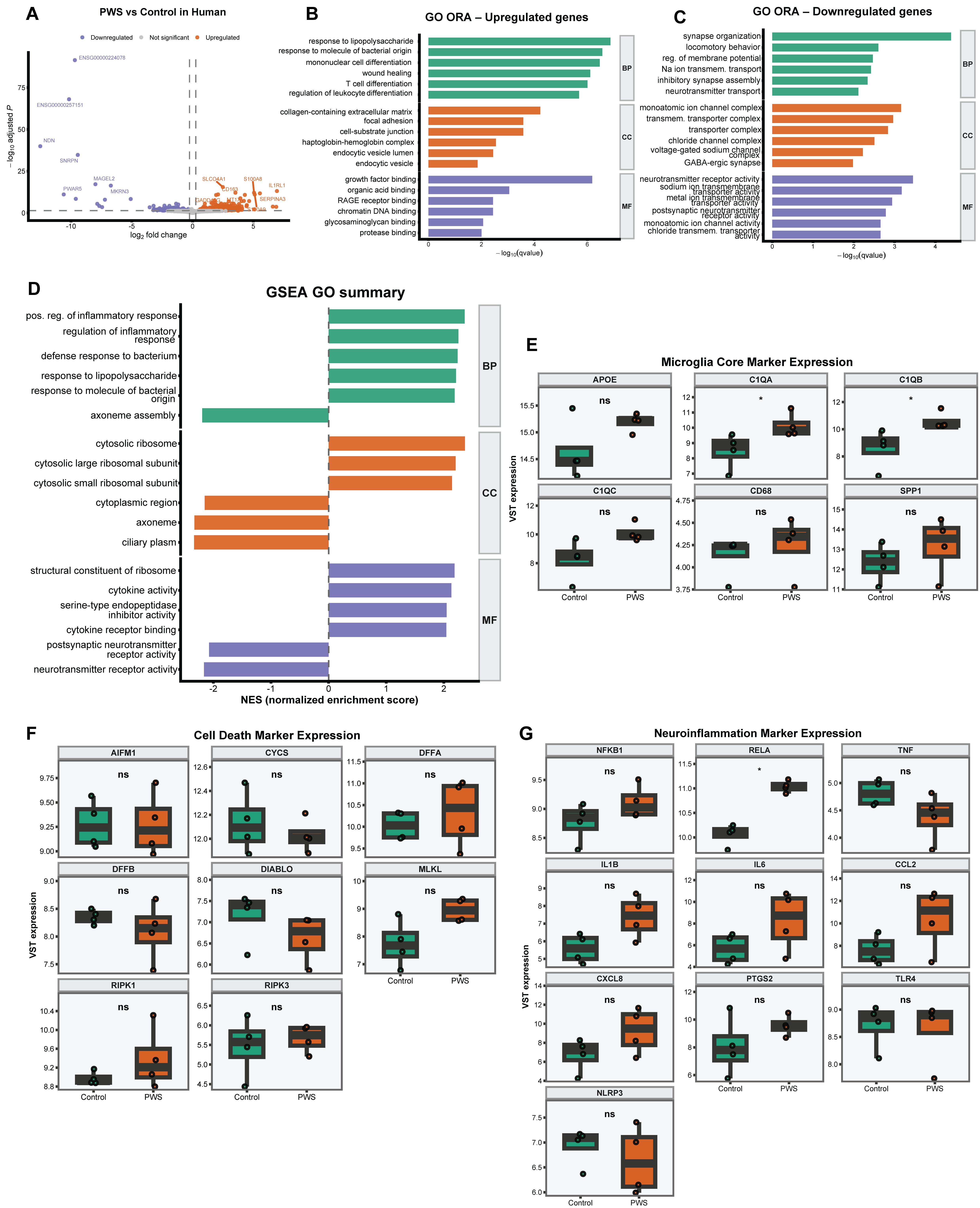

**Supplementary Figure 9**

**Bulk RNA-seq analysis of human PWS hypothalamic samples reveals immune activation and synaptic dysregulation**

**(A)** Volcano plot showing differential gene expression between Prader–Willi syndrome (PWS) and healthy control human samples. Differential expression was performed using DESeq2. Genes with FDR < 0.05 and |log2FC| ≥ 0.2 are highlighted (orange: upregulated in PWS; blue: downregulated in PWS). **(B)** Gene Ontology (GO) over-representation analysis (ORA) of upregulated genes in PWS. Enriched terms are grouped by ontology categories: Biological Process (BP), Cellular Component (CC), and Molecular Function (MF). Bar length represents statistical significance (−log10(q-value)). Enrichment highlights immune-related processes, including response to lipopolysaccharide, leukocyte differentiation, and wound healing, as well as extracellular matrix and vesicle-associated components.

**(C)** GO ORA of downregulated genes in PWS. Enriched terms are predominantly associated with synaptic organization, neurotransmitter transport, and ion channel activity, indicating impaired neuronal signaling and synaptic function. **(D)** Gene Set Enrichment Analysis (GSEA) summarizing pathway-level changes between PWS and control samples. Enriched pathways include immune and stress-related processes, alongside depletion of neuronal and synaptic signaling programs. **(E)** Expression levels of core microglial marker genes (*APOE, C1QA, C1QB, C1QC, CD68, SPP1*) across conditions, confirming the presence and transcriptional modulation of microglial signatures in bulk tissue. **(F)** Expression analysis of cell death–related genes (*AIFM1, CYCS, DFFA, DFFB, DIABLO, MLKL, RIPK1, RIPK3*), indicating altered apoptotic and survival pathways in PWS samples. **(G)** Expression of neuroinflammation-associated genes (*NFKB1, RELA, TNF, IL1B, IL6, CCL2, CXCL8, PTGS2, TLR4, NLRP3*), demonstrating increased inflammatory signaling in PWS compared to controls.
